# Disease-associated programming of cell memory in glycogen storage disorder type 1a

**DOI:** 10.1101/2023.02.20.529109

**Authors:** U Sprecher, J D’Souza, K Mishra, N Muchtar, O Shalev, A Eliassaf, A Morshina, A Canella Miliano, G Mithieux, F Rajas, S Avraham, F Castellani Moses, H Kauffman, Y Bergman, N Garti, S Garti, M Linial, Y Anikster, O Kakhlon, M Weil

## Abstract

Glycogen storage disorder type 1a (GSD1a) is caused by loss-of-function mutations in the catalytic subunit of glucose-6-phosphatase enzyme (*G6PC1*) in the liver, kidney and intestine exclusively. Here we show the surprising results that while not expressing *G6PC1*, primary skin fibroblasts isolated from GSD1a patients’ skin biopsies preserve a distinctive disease phenotype irrespective of the different culture conditions under which they grow. This discovery was initially made by phenotypic image-based high content analysis (HCA). Deeper analysis into this disease phenotype, revealed impaired lysosomal and mitochondrial functions in GSD1a cells, which were driven by a transcriptional dysregulation of the NAD^+^/NADH-Sirt-1-TFEB regulatory axis. This dysregulation impacts the normal balance between mitochondrial biogenesis and mitophagy in the patients’ cells. The distinctive GSD1a fibroblasts phenotype involves elevated H3 histone acetylation, global DNA hypomethylation, differences in the chromatin accessibility and different RNA-seq and metabolomic profiles, all of which suggesting that in some way a distinctive disease cell phenotype is programmed in these cells *in vivo* and that this phenotype is maintained *in vitro*. Supporting this notion, reversing H3 acetylation in these cells erased the original cellular phenotype in GSD1a cells. Remarkably, GHF201, an established glycogen reducing molecule, which ameliorated GSD1a pathology in a liver-targeted inducible *L.G6pc^-^* knockout mouse model, also reversed impaired cellular functions in GSD1a patients’ fibroblasts. Altogether, this experimental evidence strongly suggests that GSD1a fibroblasts express a strong and reversible disease phenotype without expressing the causal *G6PC1* gene.

## Introduction

Glycogen storage disorder type 1a (GSD1a), or von Gierke disease, is caused by deficiency in glucose-6-phosphatase (G6Pase) activity, which catalyzes the hydrolysis of glucose-6-phosphate to glucose and phosphate in the terminal steps of gluconeogenesis, and glycogenolysis-dependent glucose production in the liver, kidney, and intestine exclusively^1–5^. GSD1a is a rare disease within the group of glycogen storage disorders (GSDs) that are inherited diseases of glycogen metabolism where either the breakdown or synthesis of glycogen is disturbed. There are 19 types of GSDs^6^, and they are classified based on the specific enzyme deficiency and the affected tissue^7^. GSD1a is characterized by accumulation of glycogen and fat in the liver and kidneys, resulting in hepatomegaly, renomegaly, signs of hypoglycemia, growth failure, as well as hyperlipidemia, hypercholesterolemia, hyperlactatemia and hyperuricemia^8–11^. Currently there is no cure for GSD1a, treatment is limited to dietary interventions alone which may hold off some of the disease symptoms^11^. Since glycogen accumulation is the pathogenic factor in glycogen storage disorders, we have previously developed a cell-based assay for drug high throughput screening (HTS) to identify small molecule inhibitors of insoluble glycogen, or polyglucosan bodies (PGB) accumulation, irrespective of the mechanism of their formation, in skin fibroblasts of patients with Adult Polyglucosan Body Disease (APBD), or GSD type IV ^12^. One of these compounds, GHF201 has so far shown positive results in APBD and in GSD type III mouse models with significant improvement in their survival, metabolism and mitigation of disease progression^13,14^. Remarkably, the lysosomal protein LAMP1 was demonstrated to be the target of GHF201, improving the impaired autophagic flux, as well as mitochondrial activity in APBD patients’ fibroblasts^13^. In our efforts to evaluate GHF201 as a potential global therapeutic agent for GSDs, we opted to first characterize GSD1a patients’ primary fibroblasts phenotype as a cell model for the disease, using image-based high content analysis (HCA) as done previously for APBD fibroblasts, but having in mind that the G6Pase encoding gene, *G6PC1,* is not expressed in fibroblasts^15,16^. We hypothesized that under these circumstances GSD1a patients’ fibroblasts can serve as a useful negative control to test tissue specific effects of the novel compound. Here we show the surprising results, from phenotypic HCA experiments, that GSD1a patients’ fibroblasts have a distinctive disease phenotype as compared to healthy control (HC) fibroblasts. Key organelles such as mitochondria and lysosomes show the strongest morphometric differences in spite of the fact that *G6PC1* is not expressed in these cells. Deeper analysis into the GSD1a phenotype indicates a lower oxygen consumption rate (OCR), poor ATP production by mitochondria, and high glycolytic compensation in these cells. Also, lysosomal function and autophagy were compromised, and, expectedly, recovered by GHF201, in GSD1a fibroblasts. Furthermore, we show that PGC1α and TFEB, ^17,18^ the transcriptional regulators of mitochondrial biogenesis and lysosomal function, respectively, and the major regulators of the metabolic axis pAMPK-Sirt-1-NAD^+^/NADH are significantly inhibited in fibroblasts from GSD1a patients. Moreover, the Sirt-1-inhibitable histone H3K27 acetylation^19,20^ was elevated in the chromatin of GSD1a patients’ fibroblast and, in turn, reduced by GHF201 treatment. Importantly, the curative effect of GHF201 is not restricted to GSD1a patients’ fibroblasts and it does ameliorate established GSD1a pathology by reducing liver G6P and glycogen and increasing liver and serum glucose in a liver-targeted inducible *L.G6pc^-/-^* knockout mouse model of GSD1a. Together with this, transcriptomic and metabolomic analyses of GSD1a fibroblasts show significant differences in gene expression and key metabolites that adds up to the disease phenotype, including evidence of major deregulation of master gene regulators like Hox genes clustered in defined chromosomes, supporting a possible epigenetic regulatory change in the GSD1a cells. As we presumed that the molecular memory of the disease in GSD1a patients’ skin fibroblasts is programmed epigenetically, we thoroughly investigated epigenetic modulation mediated by both the diseased state and GHF201. In support of a general epigenetic effect, our ATAC-seq analysis indeed shows distinct differences in the chromatin accessibility between GSD1a and HC fibroblasts. Further in-depth analysis demonstrated that the DNA methylation pattern is completely modified by the diseased state. Lastly, pharmacological reversal of histone deacetylation was able to significantly mitigate phenotypic differences between GSD1a and HC cells, strongly suggesting that those differences stem from a different pattern of histone acetylation programmed by the diseased state in tissues not expressing the *G6PC1* mutant. Altogether, this experimental evidence describes a strong disease phenotype in GSD1a fibroblasts without expressing the *G6PC1* gene suggesting a disease-programmed phenotype caused by a yet not fully deciphered epigenetic mechanism. Moreover, the beneficial GHF201 effects *in vitro* and *in vivo* may consolidate further its novel therapeutic potential for GSDs.

## Results

### Phenotypic analysis of GSD1a fibroblasts by image-based HCA

To characterize the phenotype of skin fibroblasts from GSD1a patients we applied image-based HCA. For this purpose, skin fibroblasts from GSD1a patients and HC (age and gender matched) individuals were incubated in microscopy grade 96 well plates for 24 h and 48 h in cell culture media under serum and glucose starvation and for 72 h where full medium (with serum and glucose) was restored for the last 24 h of incubation. These different culture conditions were used to induce metabolic stress in the cells caused by sugar deprivation (24 and 48 h) and consecutive glycogen burden caused by adding back sugar to the media for 24 h (72 h), which proved to be highly effective in describing the GSD phenotype as shown in similar, previously published HCA phenotyping experiments of APBD (GSD-IV) and GSD III patients’ fibroblasts^13,14^. Our ultimate goal was to test whether the metabolic reprograming induced *in situ* by the patients’ diseased state before culturing generates stable epigenetic modifications withstanding seclusion from the original *in situ* environment. Thus, using the non-physiological 72 h condition, after the fibroblasts were cultured in full media remote from the *in situ* environment, can confirm the stability and environment-independence of these metabolically-driven epigenetic modulations.

At each experimental time point, the cells were simultaneously co-stained with fluorescent probes for nuclei, cytoplasm, lysosomes and active mitochondria. Live images of thousands of cells were then taken under environmentally controlled conditions using an Operetta G1 image analyzer and analyzed using Harmony 4.8 image analysis software. This procedure was repeated in three constitutive weeks to account for different passages in cells, as described in Methods. The data was subjected to several processing steps including batch effect analysis and outlier detection methods (See Methods and Figure S1a). To simplify our data presentation, we narrowed the 150 original features generated from the image analysis down to 24 using feature selection by backward feature elimination (see Methods). The selected 24 features were represented by area (size of the organelle), fluorescence intensity, and texture (combines area and intensity). To statistically confirm the phenotypic differences, two-tailed t-tests were calculated between GSD1a and HC samples for each condition (24, 48 and 72 h), and each feature (area, fluorescence intensity and texture) for each labeled organelle. To strengthen the statistical depth obtained from these experiments we applied a series of state-of-the-art statistical tools such as power analysis, Bayes factor and effect size confidence intervals (CI) (see Methods, Figure S1b and Supplementary Table 1). Figure 1a shows representative images of GSD1a and HC generated from the experiment. Box-plots of selected features representing texture of TMRE (active mitochondria), Lysotracker (lysosomes) and Calcein-AM (cytoplasm) and the area feature for Hoechst 33342 (nuclei) are shown in Figure 1b for all tested conditions. These features were arbitrarily selected from the statistical analysis mentioned above (see Supplementary Table 1) to simplify the representation of the existing phenotypic differences between the groups as shown in Figure 1. Our results indicate significant differences between GSD1a and HC samples, such as smaller nucleus areas (Hoechst 33342), lower textures of active mitochondria (TMRE) and lysosomes (Lysotracker), and higher textures of the cytoplasm (Calcein-AM) in GSD1a samples compared to HC. Scatter plots and respective effect size CI for each of the labeled organelle features were generated to confirm the statistical analysis visualization, showing the levels of the selected features in each condition (Figure 1b bottom and Figure S1b). The shade overlapping each line in Figure 1b represents a 90 percent CI which shows that there is almost no overlap between the GSD1a and HC samples in all time conditions. The observed differences between GSD1a and HC fibroblasts were confirmed by a heatmap showing the 24 selected features from this analysis, presenting the area, intensity and texture for all 4 labels in cells (Figure 1c). Figure 1c also confirms that the largest differences between HC and GSD1a cells were observed in the 72 h time condition. Principal component analysis (PCA) of the data from all the features per sample for each condition is shown in Figure 1d. The PCA shows a clear separation between HC and GSD1a fibroblasts by PC1 (55% of total variance). PC2 (24% of total variance) shows that there is also a clear separation between the different culture conditions – 24, 48 and 72 h - indicating a clear phenotypic difference between the starvation and serum restoration conditions, where GSD1a fibroblasts respond differently to the medium conditions than HC fibroblasts, (feature contribution bar plots of PC1 and PC2 are shown in Figure S1c). To further differentiate between GSD1a and HC samples and for a better insight into the main features that contribute to the phenotypic group separation, we performed linear discriminant analysis (LDA) on data from HC and GSD1a groups from all tested time conditions (Figure 1e). The LDA algorithm presented a clear separation between HC and GSD1a samples by LD1 (92% of total variance) to which features of all labeled organelles (active mitochondria, lysosomes, cytoplasm, and nuclei features) contributed. Figure S1d lists the features that contributed most to the phenotypic separation by LD1. These results show a strong image-based phenotype of the GSD1a cells in all culture conditions. Nevertheless, the 72h culture condition showed the most compelling differences between the groups. For this reason, our further study into the GSD1a phenotype shown below was done at this condition.

**Figure 1.**
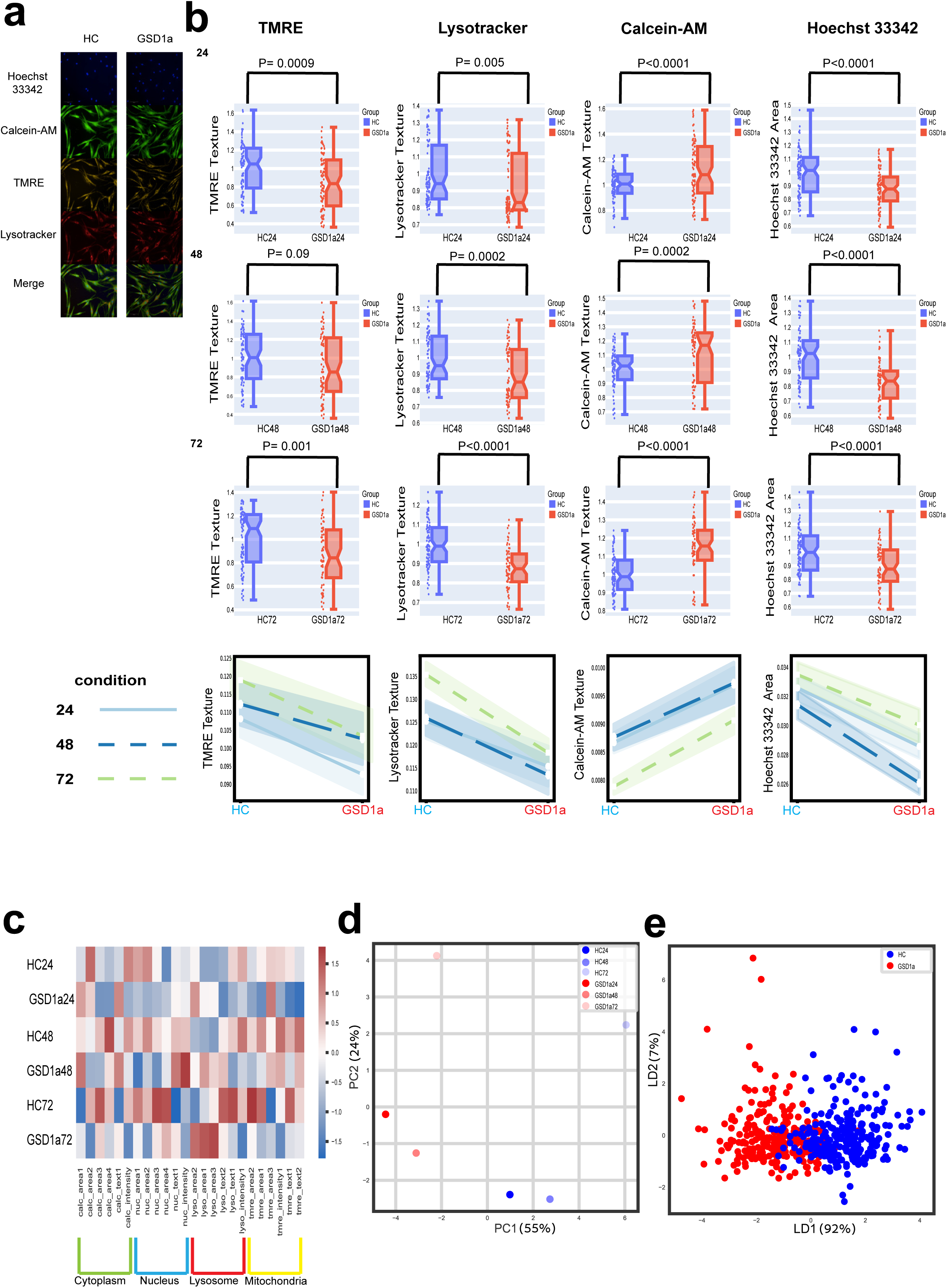
GSD1a fibroblasts image-based HCA phenotyping. **a.** Representative images of HC fibroblasts (left) and GSD1a fibroblasts (right), showcasing the organelles labeled in this experiment using Hoechst 33342 for nucleus (Blue), Calcein-AM for cytoplasm (Green), TMRE for active mitochondria (Yellow) and Lysotracker for lysosomes (Red). **b.** Up-Box plots presenting representative features comparing HC and GSD1a groups for each organelle in each time condition, from left to right-TMRE texture feature, Lysotracker texture feature, Calcein-AM texture feature and Hoechst 33342 area feature, rows show the condition from top to bottom, 24h, 48h, 72h. (N for statistical testing represents the number of wells analyzed for each group ∼ 70-85 see Supplementary table 1 for the description of each group, p values were computed using two-tailed t-tests). Bottom-scatter plots presenting the mean value for each group for each feature and each time condition tested, A 95% confidence interval computed using the data for each group is shown as a shade around the line. **c.** Heatmap representing 6 features for each organelle, each row presenting normalized average values for HC and GSD1a sample wells in each time condition. The different feature names shown for each label are numbered 1 to 6 for simplification (original feature names are mentioned in Supplementary table 1). **d.** PCA plot showing GSD1a sample wells averaged from each time condition (Red gradient), and HC sample wells averaged from each time condition (Blue gradient). **e.** LDA plot, showing all GSD1a sample wells grouped from all time conditions (Red) compared to all HC sample wells grouped from all time conditions (Blue).

### The mitochondrial function of GSD1a fibroblasts and its modulation by GHF201

To further characterize the mitochondrial and lysosomal phenotype of GSD1a fibroblasts, a deeper and functional analysis of the GSD1a mitochondrial and lysosomal phenotype was performed in patients’ skin fibroblasts in the presence or absence of our newly discovered GHF201 compound. The purpose of this set of studies is to demonstrate that, while not expressing the G6PC1 mutant, GSD1a fibroblasts present a clear bioenergetic pathology. The genuineness of this pathological phenotype is supported by its partial reversibility via lysosomal activation (by GHF201), known to correct bioenergetic aberrations. As a first step, we confirmed the effect of GHF201 on glycogen reduction and concomitant rise in intracellular glucose in GSD1a fibroblasts by Periodic acid–Schiff (PAS) assay and Mass Spectrometry (MS, these glucose data are part of our metabolomics results, Fig. 6)). The results are shown inFigure S2a. *N.B.*, glycogen levels in GSD1a fibroblasts were not significantly different than those of HC fibroblasts, probably since both do not express the *G6PC1* mutant. Next, we found that mitochondrial activity in GSD1a fibroblasts, as compared to HC, is compromised in all the parameters derived from the oxygen consumption rate (OCR) as demonstrated in our Seahorse extracellular flux experiments (Figure 2a): Basal and maximal respiration rates were lower in GSD1a cells compared with HC (Figures 2b, 2c). Furthermore, ATP production capacity, spare respiratory capacity, proton leak and non-mitochondrial respiration rates were also significantly lower in GSD1a cells compared with HC (Figures 2d-2g). GHF201 treatment significantly increased all these OCR-derived parameters (Figures 2a-2g). We hypothesized that the deficient ATP production observed in these experiments may have stimulated glycolytic compensation in GSD1a cells. To test this possibility, we performed Seahorse’s real-time ATP rate assay to simultaneously quantify ATP production from glycolysis and OxPhos. Glyco-ATP production in GSD1a cells was significantly decreased, while mito-ATP production rate was slightly increased as compared to HC, even though the OCR-predicted ATP production was decreased (Figures 2b and 2d). This apparent discrepancy demonstrates the fact that OCR only measures the capacity of the mitochondria to produce ATP, while the ATP Rate Assay provides a real-time direct measurement of ATP levels using coupling efficiency and P/O ratio assumptions. GHF201 increased the relative contribution of glycolytic ATP production at the expense of mitochondrial (OxPhos) ATP production in HC but not in GSD1a cells. Acute on-assay supplementation of GHF201 for 20 minutes was more effective at augmenting the glycolytic contribution to ATP production than 24 h (chronic) pretreatment with GHF201 in HC compared to GSD1a cells (Figure 2h). This can be explained by the extracellular acidification rate (ECAR) results showing a higher dependence on glycolysis in GSD1a cells throughout the assay (Figure 2i) indicated by higher glycolysis and glycolytic capacity (Figures 2j, 2k) with or without GHF201. To note, while glycolytic capacity in GSD1a fibroblasts is lower than that of HC fibroblasts (Figure 2h), their ECAR-based measured glycolysis and glycolytic capacity are higher. ECAR measures medium acidification and thus reflects the production of lactic acid, which is a byproduct of glycolysis. However, medium acidification is also influenced by other factors that can acidify the extracellular environment, especially CO_2_ production which can originate from the intramitochondrial Krebs cycle which produces reductive substrates for mitochondrial respiration, or OCR. Moreover, the buffering capacity of the Seahorse mito stress assay medium might mask changes in lactic acid production, leading to an underestimation of glycolytic activity. On the other hand, glycolytic ATP production measured by the ATP rate assay directly quantifies the rate of ATP production from glycolysis. Notably, there is a major difference between ECAR and the ATP rate assay: The ATP rate assay is less sensitive to variations in buffering capacity than ECAR measurements. This is because the ATP rate assay relies on inhibitor-driven changes in OCR and ECAR, rather than absolute pH values. Teleologically, the increased ECAR in Gsd1a cells represents a known compensatory response to deficient ATP production which is stimulation of glycolysis (Fig. 2i). To test the success of this known compensatory attempt, we applied the real-time ATP rate assay, which demonstrated that this compensatory response did not produce an actual increase in glycolytic ATP production in GSD1a cells as compared to HC cells. Together with this we investigated the concomitant effect of the glycolytic status of GSD1a on the mitochondrial biogenesis profile of these cells measured by the master regulator PGC1α levels in the presence and absence of GHF201. The results from these experiments show lower PGC1α levels in GSD1a cells, as compared to HC that can be increased by GHF201 treatment (Figure S3a, Figure S3d and representative images in Figure S2b). The reproducibility and validity of these results were tested by a statistical bootstrap mediated method to compute a 95% effect size CI (BCI) for each group (see Methods) which is represented in the lower panel of Figure S3a. The BCI graph demonstrates the precision of the mean difference measured between the GSD1a and GSD1a-GHF201 treated groups as compared to HC manifested by the gap between CI boundary values of each group. Following these results, we tested the protein levels of the PGC1α dependent regulators of mitochondrial biogenesis^21–23^ NRF1 and TFAM and of the positive regulator of the mitochondrial electron transfer chain cytochrome oxidase IV (COX IV) in GSD1a patients’ cells. Our Western Blot (WB) data (Figure S3b) indicate that the expression levels of NRF1, TFAM, and COX IV are significantly lower in GSD1a fibroblasts than in HC fibroblasts. Interestingly, treatment with GHF201 increased the levels of these markers.

**Figure 2.**
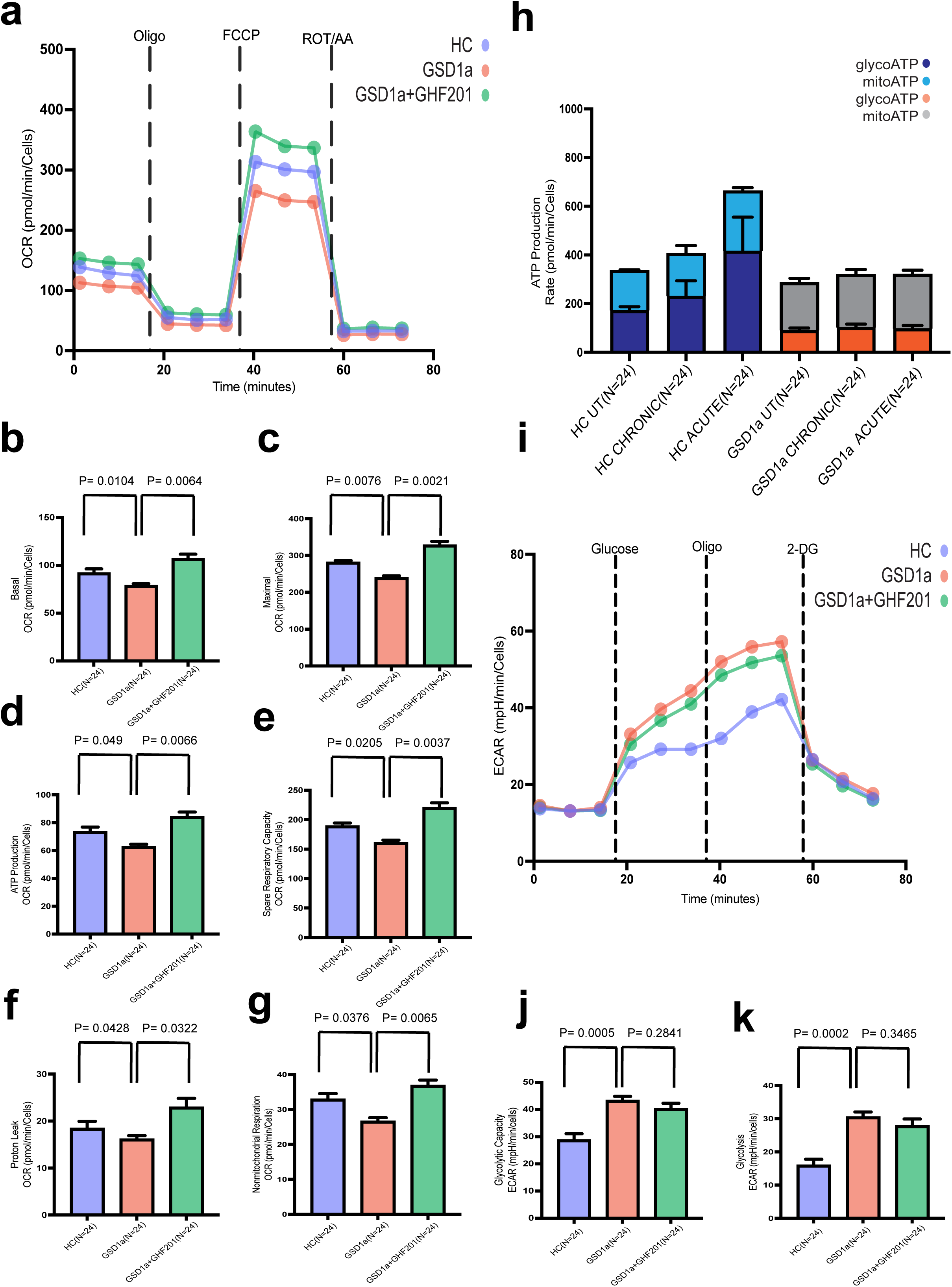
Metabolic profiling of GSD1a and HC samples treated with GHF201. **a.** Representative quantification of Seahorse real-time cellular oxygen consumption rates (OCR) of GSD1a(red), HC(blue) and GSD1a treated with 50 µM GHF201(green) fibroblasts over time. Dashed lines indicate the addition of mitochondrial inhibitors (Oligomycin; FCCP; Antimycin A/Rotenone) for assessment of mitochondrial respiration. The circle at each time point represents an average for each group at each time point. **b-g.** Changes in features derived from OCR measurements identified from GSD1a(red), HC(blue) and GSD1a treated with 50 µM GHF201(green) fibroblasts. (b – Basal respiration, c - Maximal respiration, d - ATP production, e - Spare respiratory capacity, f - Proton leak and g - Non-mitochondrial respiration(N indicates the number of wells analyzed for each group, p values were computed using two-tailed t tests). **h.** Representative quantification of real-time ATP production rates by glycolysis and OXPHOS in GSD1a, HC, HC treated with 50 µM GHF201 and GSD1a treated with 50 µM GHF201 fibroblasts. (N indicates the number of wells analyzed for each group, p values were computed using two-tailed t tests). **i.** Representative quantification of Seahorse real-time extracellular acidification rate (ECAR) of GSD1a(red), HC(blue) and GSD1a treated with 50 µM GHF201(green) fibroblasts over time. Dash line indicates the Sequential compound injections (Glucose, Oligomycin and 2-DG). The circle at each time point represents an average for each group at each time point. **j-k.** Bar graphs representing the measurements of j- glycolysis, k-glycolytic capacity from GSD1a(red), HC(blue) and GSD1a treated with 50 µM GHF201(green) fibroblasts (N indicates the number of wells analyzed for each group, p values were computed using two-tailed t tests).

### Metabolic consequences of the mitochondrial phenotype in GSD1a elucidated by the p-AMPK-NAD^+^-Sirt-1-PGC1α axis

*To place the mitochondrial phenotype of GSD1a cells in a metabolic context*, we investigated the involvement of a major metabolic regulatory pathway, p-AMPK-NAD^+^-Sirt-1-PGC1α axis^24^. To this end we performed experiments shown in Figure 3. This pathway is initiated by phosphorylation of AMPK, which was found to be low in GSD1a cells, compared to HC, as shown by immunofluorescence (IF) HCA and WB (Figure 3a, Figure S3d and representative images in Figure S2b). p-AMPK-mediated activation in turn, increases the mitochondrial respiration-dependent NAD^+^/NADH ratio, which is reduced in GSD1a (Figure 3b). The NAD^+^/NADH ratio correlates with the mitochondrial respiration rate as NADH is oxidized by complex I. This is also illustrated by our bioenergetic results, where HC cells with higher NAD^+^/NADH also manifest higher mitochondrial respiration (Figure 2). However, the biological significance of increased respiration is not necessarily an increase in ATP availability, due to the increased respiration, and ATP is not always a limiting factor for cell thriving. As stated, increased mitochondrial respiration also increases NAD^+^ availability, which plays a major role in glycolysis, TCA cycle, and amino acid, fatty acid, and nucleotide synthesis^25^. To test the relative contribution of NAD^+^ *v* ATP to cell thriving, we imposed ATP demand using the cation ionophore gramicidin, which induces ATP turnover to activate the Na/K pump and restore intracellular ion concentration gradients. Increased ATP turnover and ADP/ATP ratio by gramicidin elevated the NAD^+^/NADH ratio probably by activation of F1F0-ATPase and consequent dissipation of the mitochondrial membrane proton gradient (Figure 3c). This NAD^+^/NADH increase correlated with reduced cytotoxicity and increased cell confluence (Figure 3d) suggesting that NAD^+^ availability prevails over ATP availability as an effector of cell thriving in GSD1a cells. GHF201, both on its own and in combination with gramicidin, also increased NAD^+^/NADH ratio and cell viability, which was reversed by the respiration inhibitor oligomycin A, in further support that respiration-dependent NAD^+^ availability mediate GHF201’s corrective effect in GSD1a cells. The GHF201-mediated increase in NAD^+^ is especially interesting considering that the level of the NAD^+^-dependent histone deacetylase Sirt-1 is also elevated by the GHF201 treatment in GSD1a cells. We also found a strong difference in the levels of Sirt-1 between GSD1a and HC cells that concomitantly aligns with the above-mentioned differences in NAD^+^/NADH ratio in each of these groups (Figure 3e, Figure S3d and representative images in Figure S2b). Sirt-1 has a critical and specific epigenetic effect on the chromatin, which is deacetylation of Histone 3 at the lysine 27 acetylation site in the chromatin^19,20^. We next measured the acetylation levels in the cells and found increased acetylation of H3K27 in GSD1a as compared to HC and reversal of this acetylation pattern by GHF201 (Figure 3f). Note that in HC cells, initially showing low H2K27 acetylation, GHF201 effect was less pronounced as in GSD1a. This increased histone acetylation might be linked to the endogenous inhibition of Sirt-1 in GSD1a cells, and its reversal might be due to reactivation of Sirt-1 by the GHF201-mediated increase in NAD^+^ availability (Figure 3c). These results align well with lower levels of the mitochondrial biogenesis regulator, PGC1α, which is known to be deacetylated by Sirt-1 and phosphorylated by AMPK^26,27^, and with the positive effect on PGC1α by GHF201 in GSD1a cells (as shown in Figure S3a).

**Figure 3.**
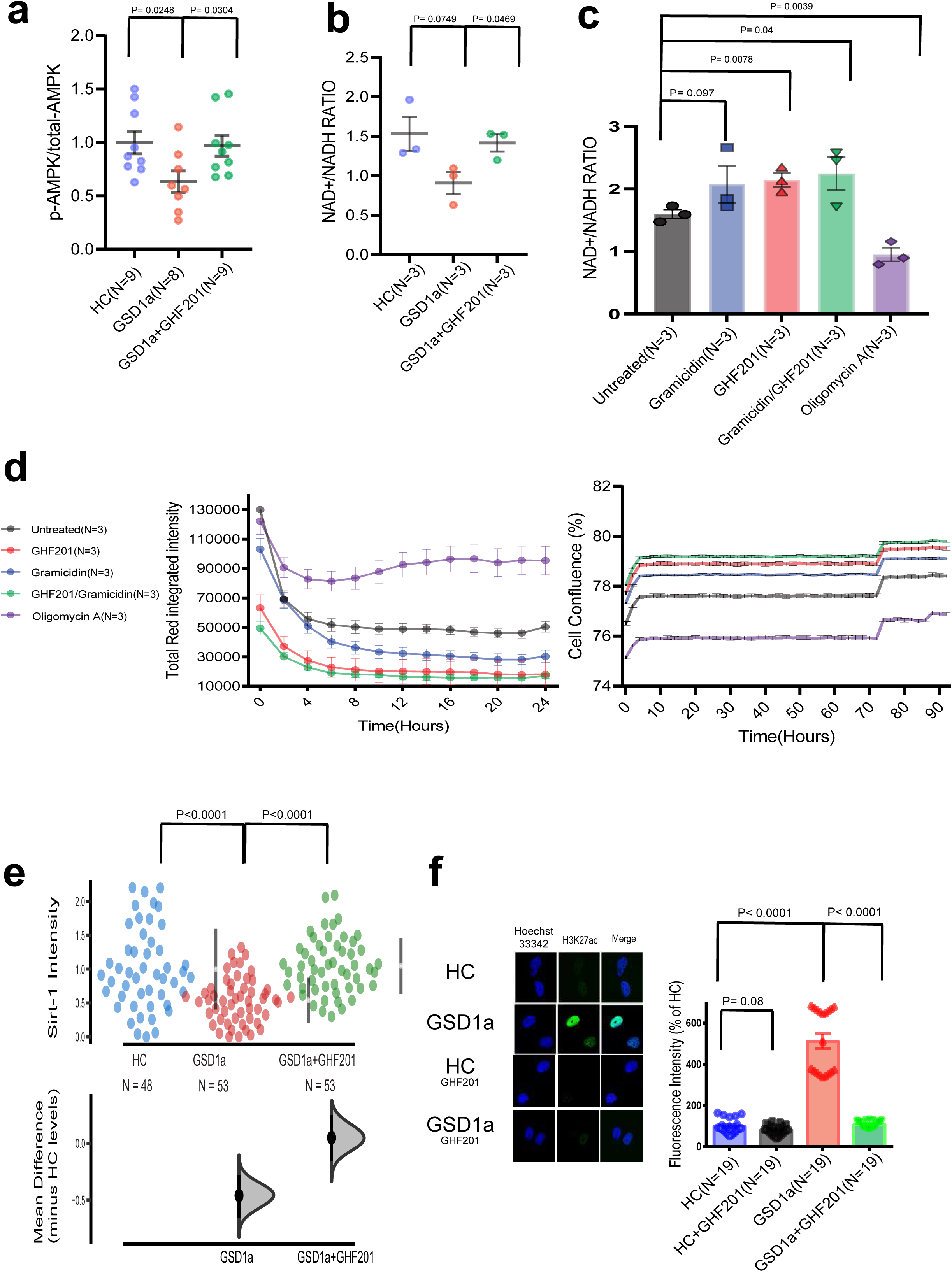
Identification of p-AMPK-NAD^+^/NADH-Sirt-1 pathway differences in GSD1a samples treated with GHF201. **a.** Scatter plots representing p-AMPK divided by total-AMPK intensity. Comparative intensity levels between HC (blue), GSD1a (red) and GSD1a treated with 50 µM GHF201 (green) fibroblasts (N indicates number of wells analyzed for each sample with p-AMPK and divided by total AMPK, p values were computed using two-tailed t tests). **b.** NAD^+^/NADH ratio measurements in HC (blue), GSD1a(red) and GSD1a treated with 50 µM GHF201(green) (N indicates number of samples analyzed for each group, p values were computed using two-tailed t tests). **c.** NAD^+^/NADH ratio measurements in GSD1a samples under different treatments-gramicidin(blue), 50 µM GHF201(red), gramicidin/GHF201 combination(green) and Oligomycin A(purple).(N indicates number of samples analyzed for each group, p values were computed using two-tailed t tests). **d.** Left-Cytotoxicity measurements in GSD1a samples over time as measured by Cytotox red and comparing different treatments. Right-Cell confluence measurements in GSD1a samples over time as measured by the Incucyte label free method comparing different treatments. Treatments included-1 µM gramicidin(blue), 50 µM GHF201(red), gramicidin/GHF201 combination(green) and 1.5 µM Oligomycin A(purple).(N indicates the number of samples analyzed for each group). **e.** Bootstrap CI Scatter plots representing Sirt-1 intensity normalized to HC levels. comparing intensity levels between HC (blue), GSD1a (red) and GSD1a treated with 50 µM GHF201(green) fibroblasts, on the right-hand side of each group a bar represents the mean intensity value with a 95% confidence interval. On the bottom the y-axis represents mean differences (effect size) between the tested groups to HC. A 95% confidence interval derived from a bootstrap resampling test is represented as a point estimate with a vertical bar and its respective distribution for each group effect size compared to HC (N indicates number of wells analyzed for each group, p values were computed using two-tailed t tests). **f.** Histone H3K27ac acetylation measurements in the nuclei of HC(blue), HC treated with 50 µM GHF201(grey), GSD1a(red) and GSD1a treated with 50 µM GHF201(green) fibroblasts. Representative nuclei images and quantification of green fluorescence in single nuclei are shown (N indicates number of wells analyzed for each group, p values were computed using One-Way ANOVA with Sidak’s correction for multiple comparisons).

### The lysosomal/mitophagic pathway in GSD1a fibroblasts and its targeting by GHF201

Having found that mitochondrial activity and biogenesis is impeded in GSD1a fibroblasts we next investigated whether malfunctioning mitochondria in GSD1a cells are poorly recycled through the process of mitophagy. To test this, we performed experiments to measure mitophagic activity^28^, which selectively removes and recycles dysfunctional and cell-damage propagating mitochondria. To this end, we used GSD1a and HC cells, stably expressing the mCherry-GFP-FIS1-leading-sequence plasmid. This includes the mCherry–GFP tag attached to the outer mitochondrial membrane localization signal of the FIS1 protein (residues 101–152), (see Methods). The results show that mitophagy is significantly impaired in the GSD1a cells demonstrated by lower mCherry/(mCherry+GFP) intensity ratio compared to HC, while GHF201 treatment manages to increase this ratio (Figure 4a). Moreover, stronger evidence for deficient autophagic regulation in GSD1a patient cells is confirmed by lower levels of the transcription factor TFEB in the cell nuclei as determined by image-based HCA using an anti-TFEB antibody (Figure 4b and representative images in Figure S2b). To confirm that the differences in lysosomal features associated with the diseased state (Figure 1) also have a functional manifestation, we measured by flow cytometry lysosomal pH using the ratiometric dye lysosensor, which quantifies pH based on the 375 nm-excited yellow/blue emission ratio. Our results (Figure 4c) show that lysosomes in GSD1a fibroblasts are more alkaline than lysosomes in HC fibroblasts. Moreover, in agreement with our previous results in APBD fibroblasts^13^, GHF201 acidified lysosomes in both HC and GSD1a fibroblasts, suggesting improvement of lysosomal function. Next, to investigate the implications of this phenotype on autophagy, we performed experiments aimed at determining the autophagic flux in GSD1a patient cells by WB analysis, as shown in Figure 4d. Compared to GSD1a cells, HC cells show higher vinblastine-dependent increase in the LC3ii/LC3i ratio (i.e., autophagic halt) indicating HC cells have higher sensitivity to this lysosomal inhibitor, or higher autophagic flux. GHF201 increased the sensitivity of the LC3ii/LC3i ratio to vinblastine indicating that it increased autophagic flux in GSD1a cells. GHF201-dependent increase in autophagic flux is also manifested by reduction in the levels of the autophagy substrate p62 (Figure S3c and d and representative images in Figure S2B).

**Figure 4.**
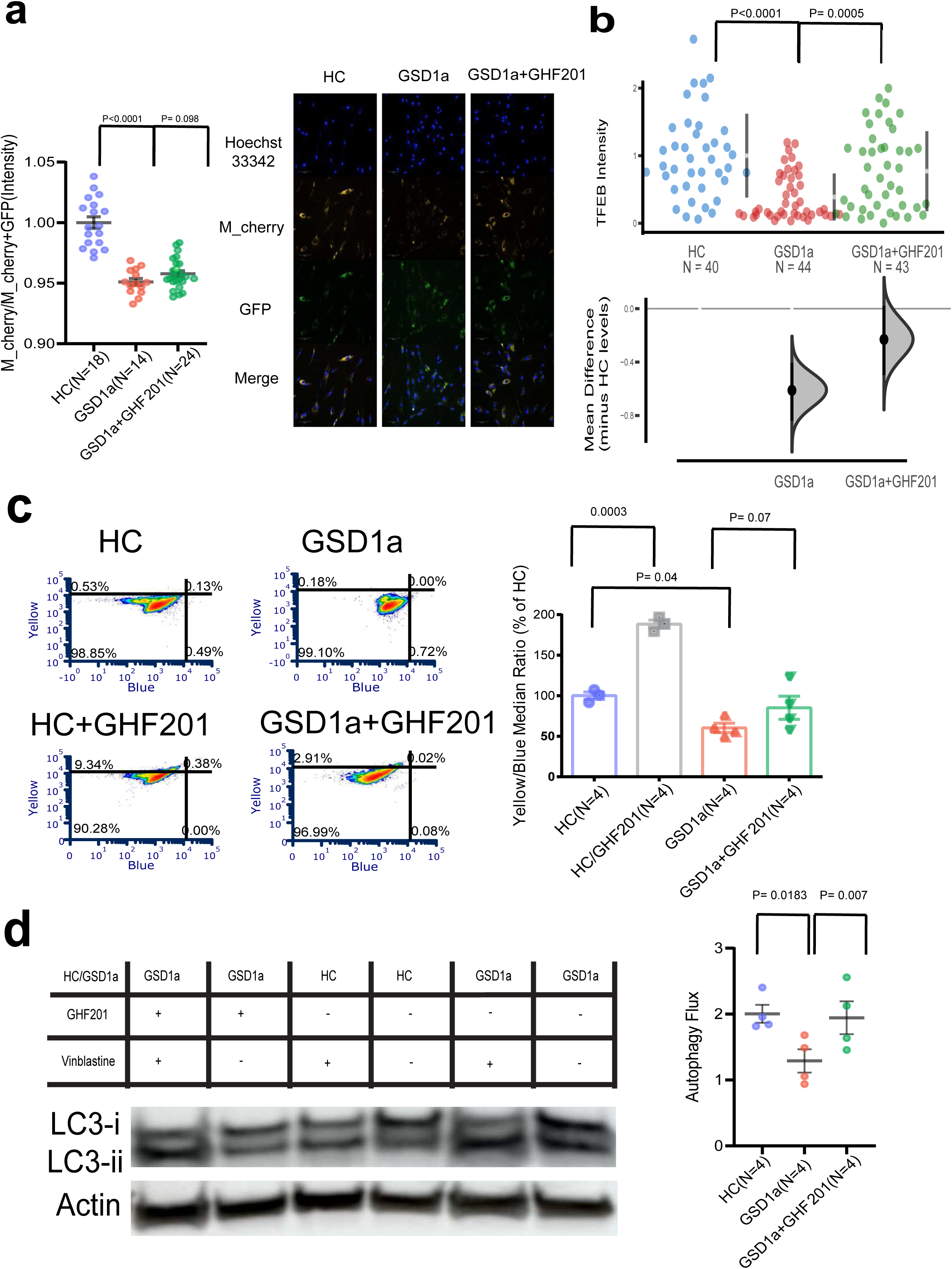
Lysosomal profiling in GSD1a and HC sample treated with GHF201. **a.** Mitophagy measurements comparisons between HC (blue), GSD1a (red) and GSD1a treated with 50 µM GHF201(green) fibroblasts stably expressing mCherry-GFP-FIS1-leading sequence, following treatment with the FCCP uncoupler. Scatter plots representing the ratio between mCherry intensity levels and the total intensity of GFP and mCherry combined, representing the extent of mitophagy, next to it shown are representative images from the experiments of all groups tested, showing from top to bottom-Hoechst 3342 (blue), mCherry (yellow), GFP (green) and a merge (N indicates number of wells analyzed for each group, p values were computed using two-tailed t tests). **b.** Bootstrap CI scatter plots representing nucleus TFEB intensity normalized to HC levels. Comparing intensity levels between HC (blue), GSD1a (red) and GSD1a treated with 50 µM GHF201(green) fibroblasts, on the right-hand side of each group a bar represents the mean intensity value with a 95% confidence interval. On the bottom the y-axis represents mean differences (effect size) between the tested groups to HC. A 95% confidence interval derived from a bootstrap resampling test is represented as a point estimate with a vertical bar and its respective distribution for each group effect size compared to HC(N indicates the number of wells analyzed for each group, p values were computed using two-tailed t tests). **c.** Lysosomal pH measurements in HC(blue), HC treated with 50 µM GHF201(grey), GSD1a(red) and GSD1a treated with 50 µM GHF201(green) as measured by flow cytometry, using the lysosomal ratiometric pH sensor Lysosensor, Left-representative examples of flow cytometry plots, Right-summarized bar graphs (N indicates samples analyzed for each group, p values were computed using two-tailed t tests). **d.** WB analysis showing autophagic flux, determined by the extent of lysosomal inhibitor (Vinblastine)-dependent increase in the ratio of lipidated to non-lipidated LC3 ratio (LC3-ii/LC3-i). Representative immunoblots and densitometric quantifications are shown comparing autophagic flux levels between HC (blue), GSD1a (red) and GSD1a treated with 50 µM GHF201(green) fibroblasts. Actin levels are shown and were used to normalize loading protein content of samples (N indicates the samples analyzed for each group, p values were computed using two-tailed t tests).

### GHF201 effect on GSD1a modeling L.G6pc^-/-^ mice

To test the *in vivo* efficacy of GHF201 on a targeted disease organ, we used the *L.G6pc^-/-^* liver-targeted inducible *G6PC1* knockout mouse model of GSD1a^29^. Following tamoxifen-induction of *G6PC1* knockout, mice were treated or not with GHF201 for 4 weeks (Figure 5a). Our results show that GHF201 treatment reduced liver G6P (Figure 5b) and glycogen (Figure 5c) and increased glucose levels in liver (Figure 5d). G6P and glucose are the main and important phenotypes of aberrant gluconeogenesis caused by G6PC1 deficiency. The reduction in G6P levels by GHF201 can be explained by enhanced lysosomal glycogen degradation with ensuing increase in cytosolic glucose emanating from this degradation. This increased cytosolic glucose might lead to enhanced glycolysis which consumes G6P. In general, G6P might also be reduced via product inhibition of hexokinase which generates it from glucose in the cytosol. However, we presume that GHF201 exerts its effect via enhancement of lysosomal glycogen degradation (the first option) similarly to the mode of action shown in previous publications, ^13,14^. Interestingly, blood glucose levels were also increased by GHF201 (Figure 5e). Hepatomegaly, or increase in liver weight in the tamoxifen-induced *L.G6pc^-/-^* mice, wasn’t corrected by GHF201 (Figure 5f). All these results point to direct therapeutic effects of GHF201 in this murine GSD1a model: GSD1a is characterized by accumulation of G6P and glycogen in the liver causing hepatomegaly and, most important clinically, hypoglycemia. All these pathogenic factors were corrected *in vivo* by GHF201 suggesting a hepato-therapeutic effect *in vivo*.

**Figure 5.**
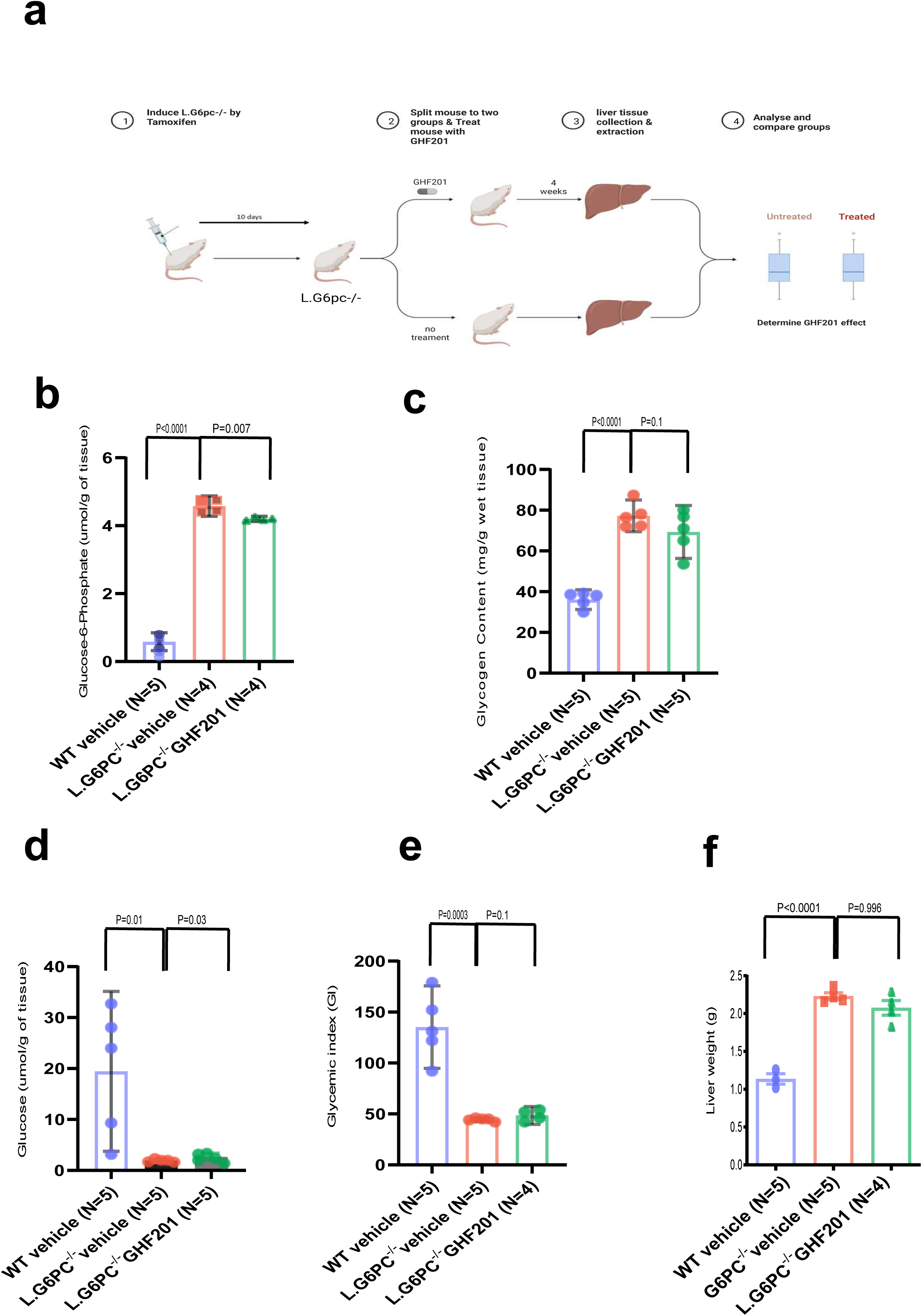
Evaluation of GHF201 in *vivo* in a LG6PC -/- GSD1a mouse model. **a.** Schematic representation of the *in vivo* experiment design for testing GHF201 on LG6PC -/- GSD1a mouse model. **b-d.** Bar plots showcasing the differences between WT vehicle(blue) LG6PC -/- vehicle(red) and LG6PC -/- GHF201 treated(green) mice in b-liver Glucose-6-phosphate, c-Liver glycogen,d-Liver glucose,e-Blood glycemic index, f-Liver weights (N indicates mice analyzed for each group, p values were computed using two-tailed t tests).

### The interaction between biochemical and cellular features

To analyze the extent of the GSD1a phenotype in patients’ primary fibroblasts and GHF201’s impact on it, we performed an integrative analysis of representative features of our multilevel phenotypic results shown in Figures 1-5 (Figure S4 and Supplementary Table 2). First, we performed a correlation analysis using the Pearson test between different representative features (Lysotracker intensity, Lysotracker area, TMRE intensity, Hoechst 33342 area, Sirt-1 intensity, autophagic flux, PGC1α intensity, TFEB intensity, P62 intensity and COX IV levels) from HCA and biochemical analysis of both the GSD1a (N=3) and HC (N=3) groups. Figure S4a shows the correlation index among all these ten features graded by a color scale. These results confirm that the phenotypic differences between GSD1a and HC, observed in lysosomes, mitochondria, and nuclei, are linked to a key biochemical marker or pathway found to be involved in the disease phenotype. For instance, autophagic flux is highly (inversely) correlated with p62 intensity (Pearson coefficient = -0.92), but not significantly correlated with nuclear (Hoechst 33342) area (Pearson coefficient = -0.12). Next, we implemented unsupervised K-means analysis on 3 HC, 3 GSD1a and 3 GSD1a treated with GHF201 samples, characterized by the relevant above mentioned biochemical data (p-AMPK/AMPK intensity, Sirt-1 intensity, NAD^+^/NADH levels, TFEB intensity, autophagic flux levels, P62 intensity, PGC1α intensity, NRF1 levels, COX IV levels, and Seahorse features; basal respiration, maximal respiration, ATP production) and tested their integration for group classification (Figure S4b). A 95% confidence interval was obtained for each group cluster. Interestingly, GHF201 treatment positions the GSD1a fibroblasts data cluster closer to the HC cluster than to the untreated GSD1a (DMSO) cluster suggesting that GHF201 treatment brings the GSD1a biochemical pathways closer to HC, which implies a curative effect. Figure S4C shows a network visualization of the interaction (based on previous knowledge^24,30–32)^ between the key markers and their representative size before and after GHF201 treatment. Overall, there are differences between the node sizes in GSD1a untreated (DMSO) and treated (GHF201) networks. Importantly, the metabolic regulator Sirt-1 pathway, dependent on NAD^+^ availability, is clearly upregulated in the presence of GHF201 which impacts on both the mitochondrial and lysosomal branches. Knowing that GHF201 targets the lysosomal marker LAMP-1 and in turn reduces glycogen accumulation (see difference in glycogen metabolite node size between groups and Figure S2a), it becomes apparent that the compound’s effect is mostly associated with the lysosomal-autophagy axis as previously demonstrated in APBD and GSD III models^13,14^. This autophagy promoting activity might directly increase the ETC rate and ATP production, which in turn increases NAD^+^/NADH ratio (Figure 3) activating Sirt-1 deacetylase activity in the GSD1a disease phenotype. This compelling evidence describes a clear GSD1a phenotype in patients’ primary fibroblast with or without GHF201 treatment and that the affected organelles found in these experiments align well with known results from other more established GSD1a models *in vivo* and *in vitro*, showing impaired mitochondrial and lysosomal pathways^3,33–36^.

### Combined RNA-seq and metabolomic phenotype characterization

To gain better understanding of the disease phenotype found in GSD1a fibroblasts we analyzed in parallel the RNA-seq and untargeted metabolomic profiles in these cells. For these experiments we analyzed RNA extracts from cells and metabolites extracted from cells and media derived from 3 GSD1a and 3 HC subjects. Results from these experiments are presented in Figure 6, Figure S5 and Figure S6. First we confirmed the low expression of *G6PC1* in our samples as shown in Figure S5a - middle. The count levels in all samples did not pass the threshold of >40 read counts per gene in our dataset. The low counts that were observed were probably a result of the depth of the sequenced reads generated in the experiment. Figure 6a (left panel) shows a PCA plot computed using the full RNA-seq dataset. A clear separation between the GSD1a and HC groups is shown based on the first component (35%) indicating a unique transcriptomic phenotype for GSD1a fibroblasts compared to HC. Differential expression analysis revealed ∼1600 differentially expressed genes (DEGs) (FDR <0.1) as shown in a volcano plot in Figure S5b (See full set of genes in Supplementary Table 3), ∼1000 genes were upregulated in the HC group compared to GSD1a and ∼600 genes were downregulated.

**Figure 6.**
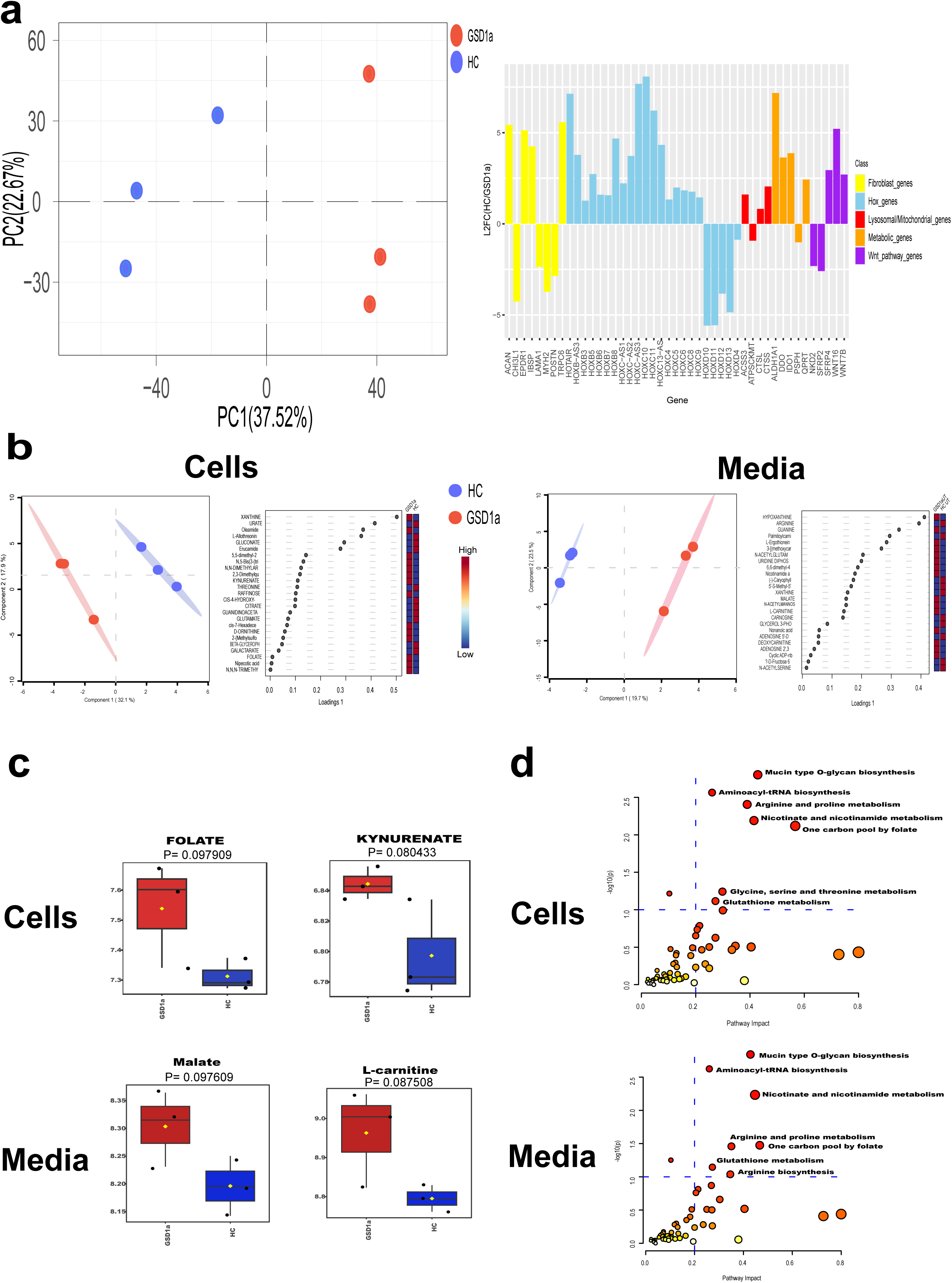
RNA-seq and Metabolomics analysis on GSD1a fibroblasts. **a.** Left-A PCA plot based on full RNA-seq data, samples represented by circles - GSD1a (red) and HC (blue). Right-A bar plot representing Log 2 Fold change ratio between HC and GSD1a samples (y axis) and selected significant genes (FDR<0.1) from differential analysis (x axis), colors of bars indicate respective cellular function for the genes. **b.** Left-sparse-PLS DA plot based on all cell metabolites, samples represented by circles - GSD1a (red) and HC (blue). Next to it, the top 25 VIP metabolites that contributed to group separation based on the first component in the sparse-PLS DA plot with their relative levels indicated by a heatmap scale (red -high, blue - low). Right-sparse-PLS DA plot based on all media metabolites, samples represented by circles - GSD1a (red) and HC (blue). Next to it, the top 25 VIP metabolites that contributed to group separation based on the first component in the sparse-PLS DA plot with their relative levels indicated by a heatmap scale (red -high, blue - low). **c.** Up-Box plots representing levels of metabolites from cell metabolomics. Down - Box plots representing levels of metabolites from media metabolomics, GSD1a in red, N=3 and HC in blue, N=3 (N indicates number of samples analyzed, p values were computed using two-tailed t tests). **d.** Up-Scatter plots from an integration analysis on significant RNA-seq genes (FDR<0.1) and metabolites from cells (p value <0.1) compared to the KEGG metabolic pathways database. Significant pathways (pathway impact score >0.2, -log10(p) >1) are indicated. Down-Scatter plots from an integration analysis on significant RNA-seq genes (FDR<0.1) and metabolites from media (P value <0.1) compared to the KEGG metabolic pathways database. Significant pathways (pathway impact score >0.2, -log10(p) >1) are indicated.

Several functionally important DEGs are presented on the right panel of Figure 6a with their log 2 fold change compared to HC and colored by their respective function in the cells: A set of Homeobox cluster genes and associated Hox expression regulators are abnormally expressed in GSD1a cells (marked in blue). HoxD4, HoxD10, HoxD11, HoxD12, HoxD13 clustered in Chromosome 2 were significantly upregulated, while HoxC4, HoxC5, HoxC6, HoxC8, HoxC9, HoxC10 and HoxC11 clustered in chromosome 12 were downregulated. Concomitantly, HoxB3, HoxB5, HoxB6, HoxB7 and HoxB8 clustered in Chromosome 17 were also downregulated. Together with this, HOXC-AS1, HoxC-AS3, HoxC-AS2, HoxC13-AS and HoxB-AS3 that are affiliated with the lncRNA class were downregulated. Another strong HoxC regulating RNA gene HOTAIR^37^, is downregulated in GSD1a fibroblasts. This gene is located within the Homeobox C (HoxC) gene cluster on chromosome 12 and is co-expressed with the HoxC genes. Among the most statistically significant differentially expressed genes in GSD1a fibroblasts are genes whose products regulate intracellular metabolic reactions, using the ESBL metabolic enzyme database^38^ we found over 60 genes overlapping with the significant differentially expressed genes in GSD1a (See full list of genes in Supplementary Table 4). Some prominent examples for metabolic regulator DEGs are IDO1, QPRT, PSPH, DDO and ALDH1A1 (a PGC1a target, marked in orange). IDO1 (Indoleamine 2,3-dioxygenase 1) and QPRT (quinolinate phosphoribosyltransferase), repressed in GSD1a patients, respectively catalyze the conversion of Trp to kynurenine and quinolinic acid to NAD^+^. Therefore, their repression corresponds to the observed decrease in cellular NAD^+^ generated by Trp metabolism as elaborated in Discussion. DDO encodes for D-Asp oxidase, a peroxisomal flavoenzyme which can degrade acidic d-amino acids (D-Asp and D-Glu) and whose decrease is therefore expected to be associated with profound intracellular metabolic changes. ALDH1A1 is also an oxidase, which oxidizes aldehydes to their corresponding carboxylic acids. The role of ALDH1A1 in detoxification of hepatic ethanol and ROS suggests that its down-modulation in GSD1a cells might be implicated in the deterioration in liver health in GSD1a. Some Wnt signaling pathway inhibitor genes,like NKD2, SFRP2 are down regulated while SFRP4 as well as Wnt Family members WNT7B and WNT16 are upregulated in GSD1a cells (marked in purple). Interestingly, downregulation of a number and not all of Wnt inhibitory genes (such as these in GSD1a cells) is associated with promoter hypermethylation in a variety of malignant glioblastomas that aberrantly increase Wnt signaling^39^. Moreover, the Wnt pathway is connected with Sirt-1 signaling, For example, Wnt signaling can decrease Sirt-1 expression and accordingly enhance p53 acetylation and activation inducing senescence in an osteoarthritis model^40^. Reciprocally, Sirt-1 can activate Wnt signaling via inhibition of the Wnt inhibitor SFRP2. This Wnt activation suppresses adipogenesis in mouse embryonic fibroblasts, committed to the adipocyte lineage^41^. Transcriptional downregulation of several genes related to the fibroblast cell type were found in GSD1a like EPDR1, IBSP and ACAN which are relevant cell adhesion molecules and extracellular matrix glycoproteins. The calcium channel TRPC6 gene related to actin stress fiber formation was also downregulated. Few other fibroblast related genes like lectin binding glycoprotein CHI3L1, myosin heavy chain MYH2 and the proteoglycan periostin POSTN and laminin LAMA1 genes were found to be significantly upregulated in GSD1a cells (marked in yellow) possibly suggesting dysregulation of extracellular matrix homeostasis. In relation to the lysosomal and mitochondrial phenotypes described above, we found several DEGs related to these metabolic organelles such as ATPSCKMT, ACSS3, CTSS and CTSL (marked in red, CTSS and CTSL are TFEB targets). ATPSCKMT was up-regulated in GSD1a fibroblasts. ATPSCKMT regulates ATP synthase activity via trimethylation of its C subunit, which enables its incorporation into the ATP synthase complex. Interestingly the up-regulation of ATPSCKMT suggests hypermethylation of a non-histone protein and activation of the mitochondrial chain. This function might serve as compensation for the mitochondrial inhibition in GSD1a (Figure 2). Deficiency in lysosomal cathepsin L protease and cathepsin S protease in GSD1a fibroblasts, on the other hand, is in line with the deficient lysosomal activity, mitophagy and mitochondrial function observed (Figures 2 and 3). Lastly, Gene Set Enrichment Analysis (GSEA) was performed on genes from the RNA-seq analysis and tested with the Gene Ontology(GO) database; Interestingly, we found changes in several pathways related to epigenetic regulation in cells as shown in a dot plot in Figure S5c. In parallel to the RNA-seq experiment, we performed an untargeted metabolomics analysis using the same samples and the same experimental design that was described for the RNA-seq experiment. Out of 183 annotated metabolites that were identified by MS in the cell extracts, approximately 30 significantly different metabolites were found comparing the cell extracts of GSD1a and HC samples. On the other hand, out of 217 annotated metabolites identified in the cell media extracts, 20 metabolites were discovered, which were significantly different between GSD1a and HC samples (p Value <0.1, see volcano plots in Figure S6b, and tables of significant metabolites in cells and media in Supplementary Table 5 and Supplementary Table 6, respectively). Figure 6b shows a sparse-PLS-DA analysis plot that was performed on both cells and media metabolomics datasets to define metabolites significantly contributing to the HC-GSD1a classification. The sparse-PLS-DA analysis revealed a clear separation by the first component (cells -32%, media-20%) between groups for both cell (Left) and media (Right) metabolomes. Next to each sparse-PLS-DA analysis plot shown are VIP (metabolites importance score) plots with the top 25 metabolites contributing to the variation explained by the first component. Prominent metabolites modified are shown in Figure 6c: In patient cells (upper panel), folic and kynurenic acids accumulate, while in patient cell media (lower panel), L-carnitine and malate accumulate. Next, we performed Metabolite Set Enrichment Analysis (MSEA) on the data, MSEA showed several metabolic pathways enriched in GSD1a samples as compared to HC. Two notable enriched pathways in both cells and media are the malate-aspartate shuttle, and the valine, leucine and isoleucine (branched chain amino acids) degradation pathways. The enrichment of the malate-aspartate shuttle was deduced from declines in glutamate, malate, L-Asp, and oxoglutarate (α-ketoglutarate) and so was inhibited as mentioned above. Enrichment of branched chain amino acid degradation was deduced by changes in Ile, Leu, Glu, pyridoxal, α-ketoglutarate, succinate, and L-Val. The decrease in the levels of the products of this pathway Glu, α-ketoglutarate, succinate, and L-Val suggests that branched chain amino acid degradation was down modulated in GSD1a fibroblasts as compared to HC fibroblasts. This conclusion is supported by the accumulation of Leu which feeds Acetyl CoA directly to the Krebs cycle, bypassing pyruvate dehydrogenase and is thus a major determinant of cellular energetic plasticity. MSEA plots for cells and media are shown in Figure S6C. To gain deeper insights regarding the phenotypic relevance of these large datasets to the GSD1a disease phenotype we integrated the RNA-seq and metabolomics datasets and tested them against the KEGG metabolic pathway database using the full set of significant differentially expressed genes from the RNA-seq experiment (FDR<0.1) and the significant metabolites from cells and media (p value <0.1). Scatter plots of the enriched pathways (p Value <0.1, pathway impact score >0.2) are shown in Figure 6d (cells, upper panel; media, lower panel). Few interesting metabolic pathway enrichments were observed. Among them, inhibition in folate and single carbon metabolism (one carbon pool by folate) and in Trp metabolism (or nicotinate and nicotinamide metabolism, which are part of Trp metabolism) in both cells and media. These pathways might be significantly implicated in hypomethylation and NAD^+^ depletion, respectively, as detailed in Discussion.

### Exploring epigenetics as a phenotypic driver in GSD1a fibroblasts by ATAC-seq analysis

Following all the above results we hypothesized that the phenotype depth described in GSD1a fibroblasts is a result of an epigenetic effect, presumably caused by the aberrant metabolism in GSD1a fibroblasts, which sustains the phenotype without the expression of *G6PC1* in these cells. To investigate this possibility, we performed ATAC-seq analysis on fibroblast samples from 2 GSD1a patients and 2 matching HC (Figure 7 and Figure S7). To explore the accessible genomic regions in GSD1a, compared to HC fibroblasts, we performed an occupancy-based analysis which can be used to identify group-unique accessible sites in the chromatin, Figure 7a shows a PCA plot results of occupancy-based analysis, which clusters GSD1a fibroblasts by PC1 (63%) and separates them from HC fibroblasts. We then performed affinity analysis using DiffBind and comparing GSD1a to HC fibroblasts (see full table of significant differentially accessible peaks in Supplementary Table 7). The heatmap (Figure 7a right) presents clear differences between the tested groups as indicated by the color scale of normalized reads of binding sites, green and red representing low and high affinity of binding sites respectively. To investigate the pattern shown in these different binding sites between the groups we computed log2 of normalized reads for each group and plotted them using box plots. This analysis revealed that the pattern of the significantly different binding sites was upregulated or downregulated in GSD1a fibroblasts as compared to HC (Figure S7b). Finally, an MA plot, that shows the log fold-change between the two groups vs mean read count, presents the differential sites found in this analysis, and corroborates the results shown in the box plots which indicate that the pattern of the differential binding sites is both up regulated (456 peaks) and down regulated (500 peaks) in HC compared to GSD1a fibroblasts (Figure 7b left). To Further investigate specific genes that were mapped to differentially accessible peaks, we plotted the peaks that represent the SLC2A8, HoxD11, HAS1, ARG1, MAP1LC3B2 and A4GALT genes. These genes show relevance for GSD1a pathology and unique phenotypes: Open chromatin in the HoxD11 and ARG1 loci, suggesting their up-regulation, is respectively associated with activation of developmental processes and with glucose-6-phosphate accumulation through gluconeogenic metabolism of fumarate released from the urea (ornithine) cycle driven by ARG1. In agreement with its open chromatin, HoxD11 expression was indeed increased. For ARG1, while its RNA read counts are below the threshold in the patient fibroblasts, possibly since ARG1 does not have a functional role in skin cells, its epigenetic activation, suggested by the open chromatin in its promoter region, might represent a relic of an adaptive activation of gluconeogenesis, vis a vis its innate dysfunction in GSD1a, in early-stage pluripotent embryonal tissue. Closed chromatin with ensuing down modulation of expression (Figure S7c), is observed in the loci of HAS1, SLC2A8, A4GALT, and MAP1LC3B2. HAS1 inhibition is associated with reduction in biosynthesis of the glycosaminoglycan hyaluronic acid, which correlates with glycogen accumulation and ensuing reduction in glucosamine and glycosaminoglycans biosynthesis^42^. SLC2A8 inhibition is associated with reduced glucose uptake, possibly as a compensatory response which would balance the potential increase in glycogen synthesis observed in GSD1a cells expressing the G6PC1 mutant. SLC2A8 inhibition can illustrate a possibly metabolically-driven systemic epigenetic change aimed at counteracting glycogen accumulation in all cells, which would actually reduce glycogen accumulation in GSD1a cell types that express the G6PC1 mutant. A4GALT inhibition leads to galactose-induced phosphate depletion and subsequent inhibition of glycogen breakdown by the phosphate-dependent glycogen phosphorylase^43^. Lastly, the closed MAP1LC3B2 locus is associated with the observed inhibited autophagy and mitophagy in GSD1a cells (Figure 7b right). To elucidate the genes and pathways that are affected by these differential binding sites, we performed GSEA using the GO database. Results from this analysis are represented in a net plot graph in Figure 7c where significantly enriched pathways, their interactions and the number of genes contributing to each pathway are indicated. From this analysis we found pathways that are related to the phenotypic characteristics identified above (see Figures 1-5) such as autophagy, mitochondrial dynamics, protein catabolic process and lysosome as well as epigenetic related pathways such as chromatin binding. Next, to assess DNA binding transcription factors that might interact with the differential chromatin areas found and affect gene expression, we performed motif analysis on the significant peaks generated from this experiment. Figure 7d presents 5 Motifs chosen from the motif analysis on up-regulated peaks in GSD1a (binding sites for FRA2, NRF2, ETS1, HOXA3, HIF1A) and 5 Motifs chosen from the motif analysis on down-regulated peaks in GSD1a (binding sites for AP-1, ETV4, KLF17, NANOG, HOXC6). The expression of the TFs associated with these motifs was confirmed as shown in a heatmap in Figure S7c. Interestingly, these TFs regulate genes involved in embryonic development and morphogenesis (HOXA3 and HOXC6), but mostly of metabolic genes: FRA-2 is a component of the AP-1 complex, which regulates, among other functions, glycolysis and cholesterol synthesis^44^. NRF2 is a key regulator of antioxidases such as glutathione peroxidases^45,46^. ETS1 and its variant ETV4 are also inducers of antioxidant enzymes and inhibitors of OxPhos activators^47,48^ and HIF1A is the key enzyme enabling the cells to cope with hypoxic stress. Notably, the down-modulated TF KLF17 belongs to the Kruppel-like TF, which directly regulate metabolic homeostasis^49^. NANOG is a key regulator of OxPhos and fatty acid metabolism^50^. Altogether, these results strongly support that a broad epigenetic difference exists between GSD1a and HC fibroblasts both at the chromatin level, by differential binding accessibility and these, accompanied by selective gene expression levels of some genes shown to be differentially expressed in GSD1a fibroblasts. The high proportion of metabolic genes driven by these transcriptional regulators suggests that the metabolic aberrations caused by the diseased state in GSD1a patients generates a metabolic feedback response in the cells.

**Figure 7.**
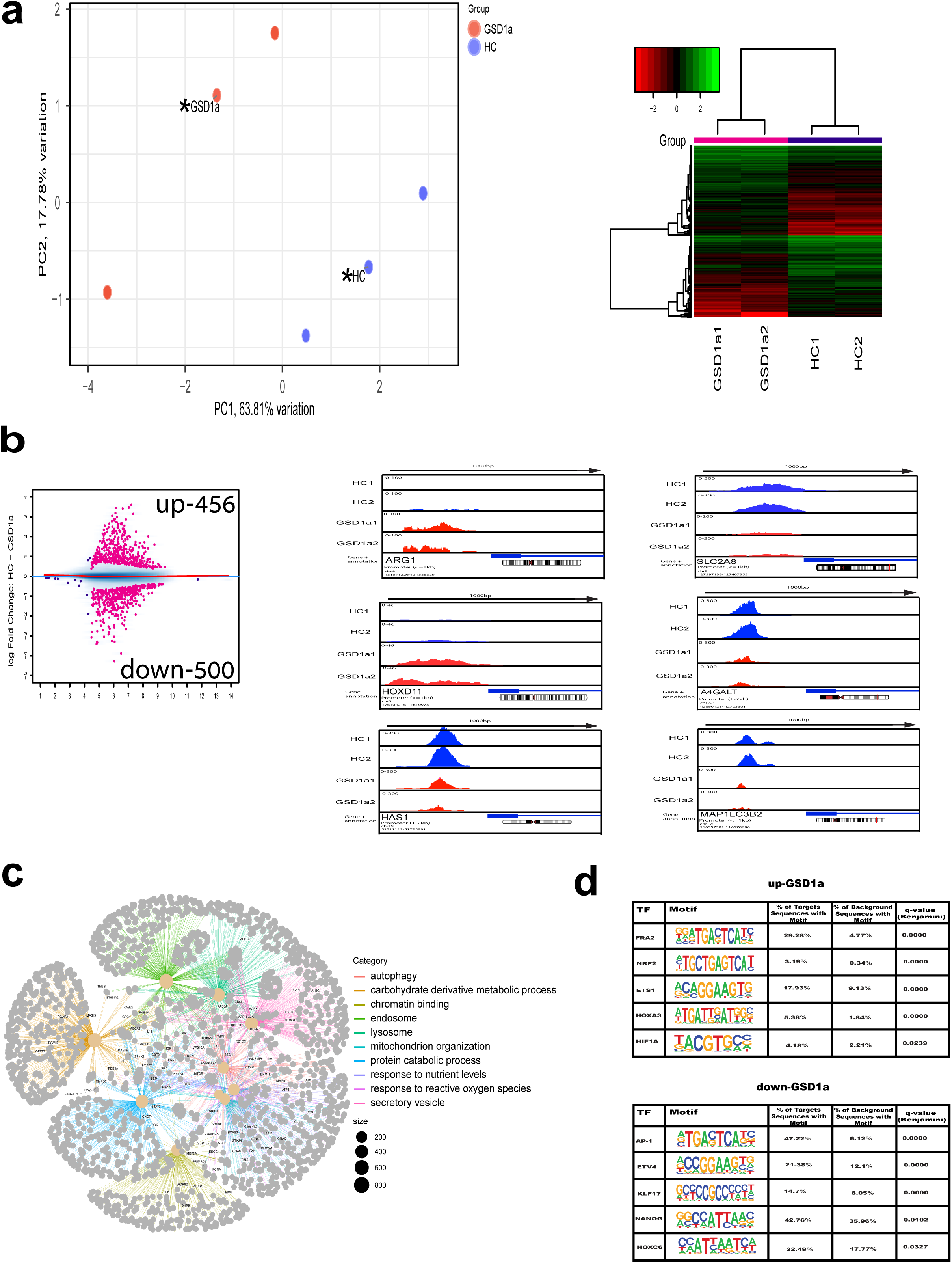
ATAC-seq analysis of global chromatin accessibility in GSD1a fibroblasts. **a.** Left-A PCA plot showing results of an occupancy analysis. Indexed samples representing the samples analyzed GSD1a (red) and HC (blue) while the un-indexed dot (black asterisk) presents the combined peaks between group samples. Right-A heatmap showing the results from the affinity analysis performed on 2 GSD1a and 2 HC samples (green: high affinity; red: low affinity, FDR<0.1). **b.** Left-An MA plot showing log 2 fold change differences between HC samples compared to GSD1a samples, the up and down regulated significant binding sites are marked in pink (FDR<0.1). Right-IGV peak plots of genes selected from the significant differentially expressed peaks(FDR<0.1), red peaks represent GSD1a samples and blue peaks represent HC samples. On the bottom of each gene plot, the respective chromosome is shown in addition to the location of the gene which is noted by a red line. **c.** A net plot presenting results from a GSEA comparing GSD1a and HC genes and integrating relevant pathways. Node size reflects gene set size and color of clusters indicate different pathways identified (FDR<0.1). **d.** Tables of selected significant motifs from ATAC-seq peak data, Tfs, Motifs with their statistics are shown. In the upper panel, motifs generated from GSD1a upregulated peaks and on the bottom motifs generated from GSD1a downregulated peaks.

### Differential methylation pattern in GSD1a fibroblasts

Following the ATAC-seq results indicating clear phenotypic differences in the open chromatin state between GSD1a and HC cells we investigated first the global pattern of non-methylated regions of genomic DNA in the GSD1a and HC groups as determined by Simultaneous Global Labeling (SIGL, see Methods). These results show reduced unmethylated CpG regions in HC cells indicating relatively lower levels of DNA methylation in GSD1a fibroblasts (see Figure S8a). To investigate in depth this differential DNA methylation profile, we analyzed differences in the genomic methylation pattern between 4 GSD1a and 4 HC samples using the Illumina EPIC V2 array. This array identifies over 900K CpG methylation probes representing important methylation sites across the human genome. Figure 8 and Figure S8 show the results of these experiments. To assess the global differences between samples from the two groups, we used the full dataset of processed probes and initially performed a regression analysis with the aim to assess the in-group variability for each of HC and GSD1a groups tested. Results from this analysis showed lower variability in the GSD1a group which implies that GSD1a fibroblasts present a methylation phenotype that is consistent between group samples as compared to the HC samples (See Figure S8b). These differences are clearly observed by an unsupervised clustering analysis that we performed based on K-means searching for two clusters between our 8 tested samples and by PCA, as shown in Figure 8a. The left panel shows the results of the clustering analysis as a cluster map. All 4 HC samples were clustered in cluster 2, while GSD1a samples were clustered in cluster 1. The PCA analysis, on the right panel of Figure 8a, shows a clear separation between GSD1a and HC cells by the two top principal components (PC1 30%, PC2 21%), In addition, the 95% CI ellipsoids that were computed for each group are clearly different in size, where GSD1a cells are within a much smaller area as compared to the HC area, indicating less sample variation within the disease group. Next, differential methylation analysis comparing the GSD1a to HC cells was performed. Figure 8b presents thousands of significantly different probe signals between the groups as illustrated by a volcano plot, on the left panel. The four top significant probes from this analysis are presented as scatter plots on the right panel (see full table of significant differential probes in Supplementary Table 8). These results show strong differences in many of the methylation sites analyzed between the groups, further strengthening the conclusion that the methylation profile of GSD1a cells is different than that of HC cells. We next performed differential methylation region analysis which is based on methylated probes distribution located at similar regions in the genome. We obtained many differentially methylated regions (DMR) between the groups (see full table of significant differential regions in Supplementary Table 9). The most significant region is presented in Figure 8C. This region is in chromosome 12, where we have revealed downregulation of the HoxC gene cluster expression (Figure 5), and where strong differences in the methylation pattern between GSD1a and HC samples are clearly visible across the chromosome’s regions. In addition to the most significant DMR (DMR 1), Figure 8c also shows two other DMRs (DMR52, DMR20981) in which GSD1a and HC samples have a different methylation pattern. At the bottom of the plot, the scale bar represents the methylation levels of the probes. The heatmap bars above it show the enhanced variability among HC samples compared to GSD1a samples. The line under the heatmap bars represents the mean methylation value of GSD1a and HC fibroblasts, indicating a strong difference in methylation profile between GSD1a and HC cells. Figure 8d shows GSEA results in a bar plot comparing GSD1a cells to HC cells using the methylation dataset and the GO database. Among the significantly enriched pathways found are lysosomal and metabolic pathways, strengthening the phenotypic differences observed previously (Figures 1-5). In addition, pathways related to epigenetic modifications are also differentially enriched between GSD1a and HC cells, implying an epigenetic origin of the GSD1a phenotype in patients’ fibroblasts cells.

**Figure 8.**
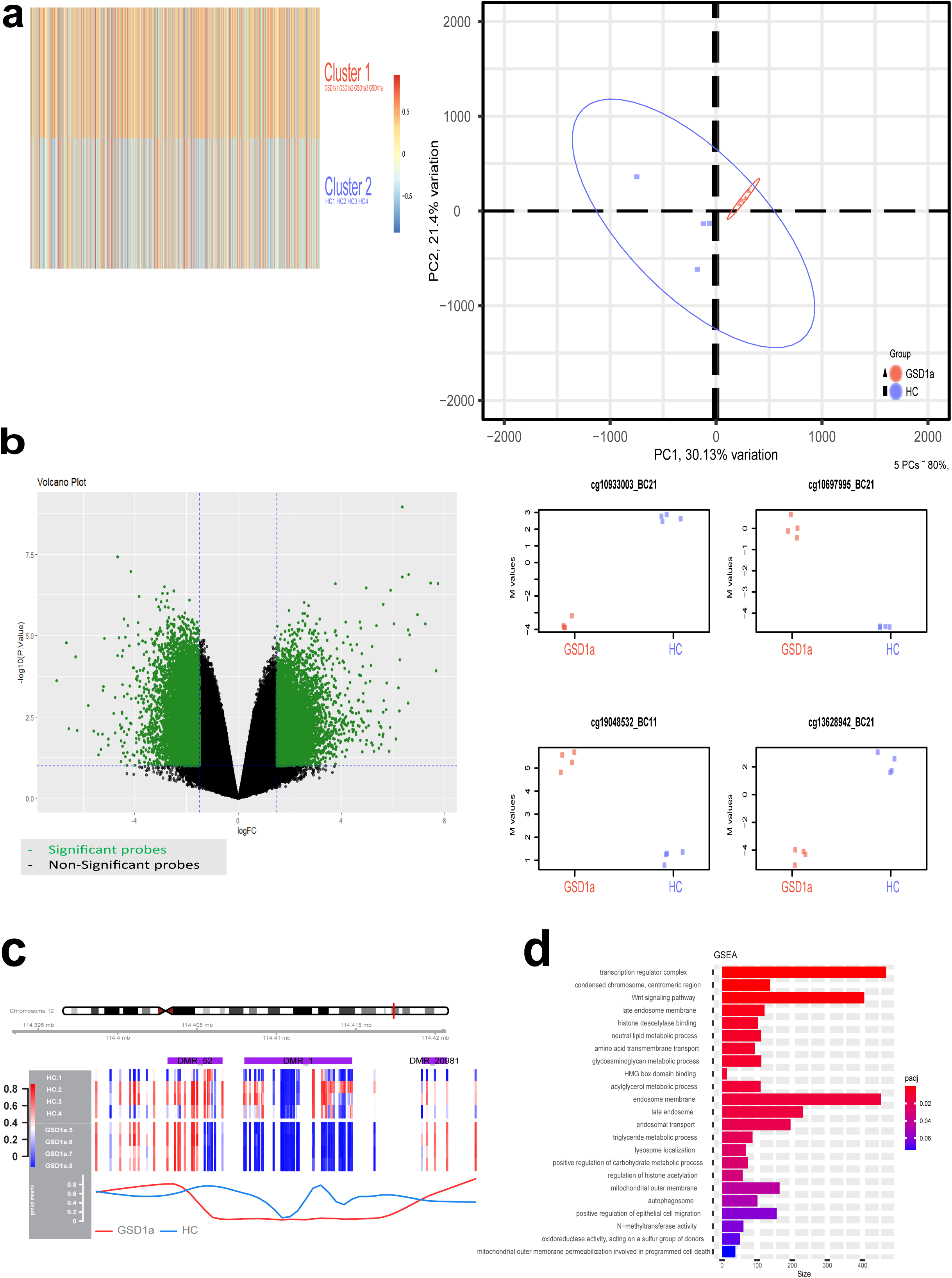
Analysis of EPIC methylation pattern in GSD1a fibroblasts. **a.** Left-a cluster map of K-means clustering results of the full EPIC methylation dataset. Right-a PCA plot based on the full EPIC methylation dataset, samples are represented by - HC (squares-blue) and GSD1a (triangles- red), groups are encircled by a 95 % confidence interval, indicating their sample heterogeneity proportional to the size of the circle. **b.** Left-volcano plot presenting results from differential methylation analysis comparing HC and GSD1a cells, green dots represent significant probes(FDR<0.1). Right-scatter plots presenting the top four significant probes computed from the differential methylation analysis and comparing GSD1a in red, N=4 and HC in blue, N=4 (N indicates number of samples analyzed, FDR<0.1). **c.** A DMR plot representing a differential methylation region analysis of Chromosome 12 comparing methylated regions between GSD1a and HC cells. The location on the chromosome of the region of interest is indicated by a red bar. The color scaled bar on the left, indicates methylation levels of GSD1a and HC samples and the red and blue lines at the bottom represents the mean methylation values for the GSD1a(N=4) and HC(N=4) groups respectively (N indicates number of samples analyzed, FDR<0.1). **d.** Bar plots of a GSEA performed on genes annotated to the methylation probes identified from the differential methylation analysis between the groups (FDR<0.1).

### ATAC-seq, RNA-seq and EPIC methylation data integration

To gain a deeper insight on the epigenetic phenotype identified in GSD1a cells (Figures 6, 7 and 8), we integrated the ATAC-seq, RNA-seq and EPIC methylation data sets. Results from this analysis are shown in Figure 9 and Figure S9. First, we examined the distribution of the chromosomes which represented the significant genes in each of the datasets (FDR<0.1). Pie charts of the chromosome distribution of each dataset are shown in Figure S9a. We found consistency between datasets with dominance of significant genes from chr1, chr2, chr3, chr7 and chr12. Next, we performed an overlap analysis on significant genes from each dataset as shown in Venn diagrams in Figure 9a, underneath each Venn diagram we plotted the regulation (direction of fold change for both datasets) of the overlapping genes (full list of overlapping genes from this analysis are shown in Supplementary Table 10). First, we found 48 overlapping genes between ATAC-seq and EPIC methylation datasets. Annotation of the genomic region of the overlapping peaks is shown in Figure S9b left-bottom. Among these 48 genes we identified several genes related to the metabolic phenotype described above such as ATP2C1, HK1, GRAMD1B, LRRC2 and SLC36A3. Next, we found 77 overlapping genes between ATAC-seq and RNA-seq datasets. Annotation of the genomic region of the overlapping peaks is shown in Figure S9b right-bottom. Here we identified several genes related to the Hox genes mentioned above; HoxD11, HoxD12 and HoxD13, HoxB5, and HoxC4. In addition, we found many genes related to metabolic pathways such as SLC2A8, GXYLT2, GCNT3 and ATP2B1. Lastly, we also identified several genes related to chromatin regulation such as NFIB and SMYD3. Finally, we found 72 overlapping genes between the RNA-seq and EPIC methylation datasets, among them several metabolism related genes such as ALDH1L2, ATP13A4, CES3 and HSD11B1, in addition to several transcription factors - KLF13, L3MBTL4 and ZBTB38. To validate the significance of the overlapping genes found between the datasets we performed permutation testing on each overlap analysis as shown in histograms in Figure S9b top. This analysis showed that the overlaps identified between all datasets were significant (p value = 0). To investigate significant genes across all datasets we extracted the overlapping genes from all three datasets as shown in a Venn diagram in Figure 9b. Regulation of the overlapping genes is shown below the Venn diagram. Results from this analysis revealed 9 shared genes (BCKDHB, HLX, ITGA11, LGI1, MBNL2, PCSK5, PTPRO, USH2A, ZBTB16). Interestingly, enrichment analysis using text mining of associations between genes and human diseases (the Jensen Diseases database) shows that all these genes are associated with carcinoma, the only disease out of 43 diseases text-mined to be associated with these 9 genes, in which all 9 genes are clustered (Figure 9c). The odds ratio for such an association of all genes was 78,138, more than two orders of magnitude higher than the odds ratio of the association between other diseases and genes from that list. Hepatocellular carcinoma is a well-established co-morbidity of GSD1a, usually presenting at later stages of the disease. The fact that carcinoma was the only disease associated with all significant genes overlapping among ATAC-seq, RNA-seq and EPIC methylation analyses indicates that there might be an etiological link between the expression of these genes and GSD1a pathology. While the patients from whom the fibroblasts were derived did not present with hepatocellular carcinoma, they were all adolescents and may therefore be predisposed to develop this co-morbidity later on. To further explore the relationship between RNA-seq, ATAC-seq promoter peaks and EPIC methylation data we integrated data of 2 HC and 2 GSD1a samples from all data sets. Then using genes which were shared across datasets we performed sparse-PLS-DA analysis and found a clear separation between HC and GSD1a samples as shown in Figure 9d. The contribution of each data set to this analysis and genes which were most relevant to group separation are shown in Figure S9c. To draw more insights from these complex datasets and taking into consideration the limited sample size we have, we implemented a bootstrap mediated approach. A bootstrap correlation analysis was performed on the full data set of the RNA-seq, EPIC methylation and on promoter regions from ATAC-seq. Using the mean correlation value generated from the bootstrap analysis we subsetted genes based on high correlation (Pearson coefficient >0.7, Pearson coefficient confidence interval < 1) and then filtered them for genes which showed high correlation between RNA-seq and ATAC-seq promoter peaks and low correlation with EPIC methylation (see Methods). Using the ∼200 genes that passed the threshold, we analyzed them using STRING^51^ (see full data set in Supplementary table 11). We then applied K-means clustering on the network of genes which resulted in 9 clusters based on centroids^51^. 7 clusters from this analysis are shown in Figure S9c and pathway enrichment analysis results from each cluster are shown in a table in Figure 9e. Results from this analysis showed many relevant pathways that align with GSD1a pathology, such as Branched chain amino acid (cluster1), Mitochondrial Fatty Acid Beta-Oxidation (cluster3), Lactate dehydrogenase activity, and Malic oxidoreductase (cluster5) and Nonalcoholic fatty liver disease (cluster6). In addition, we found several pathways that corroborate our findings from Figures 2 and 6 such as Activation of anterior HOX genes in hindbrain development during early embryogenesis (cluster2), Cadherin C-terminal cytoplasmic tail, Catenin-binding region (cluster7) and Mixed, incl. 2,4-dienoyl-CoA reductase (NADPH) activity, and ATP export (cluster4). Altogether, these results support the epigenetic phenotype found in GSD1a cells and explains more profoundly the phenotypic characteristics of GSD1a fibroblasts.

**Figure 9.**
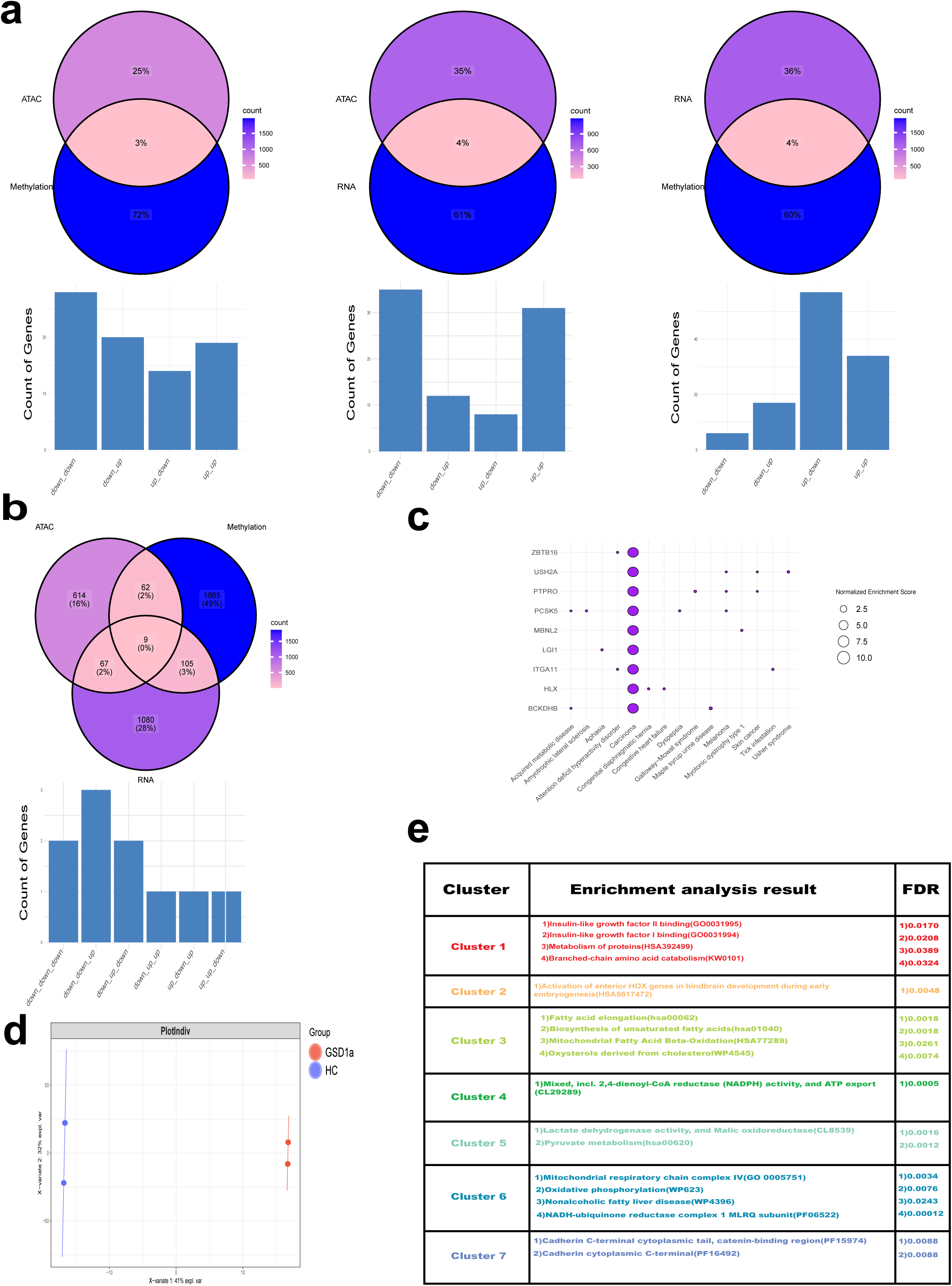
Omics integration. **a.** Venn diagrams of overlapping significant genes (FDR<0.1) between ATAC-seq and EPIC methylation data(left), ATAC-seq and RNA-seq data(middle) and EPIC methylation and RNA-seq data(right), percentages and the color scale reflect genes in each data set, underneath each venn diagram shown are bar plots indicating the pattern of log 2 fold change of the overlapping genes. **b.** Venn diagrams of overlapping significant genes (FDR<0.1) between ATAC-seq and EPIC methylation and RNA-seq data, underneath shown is a bar plot indicating the pattern of log 2 fold change of the overlapping genes. **c.** A dot plot of enriched pathways based on enrichment analysis performed on 9 overlapping genes from ATAC-seq and EPIC methylation and RNA-seq data (shown on y axis), the top 15 pathways (FDR<0.05) are shown on the x axis, purple circles indicate genes that contributed to enrichment of each pathway and the size of the circles indicate the normalized enrichment score. **d.** Sparse-PLS DA plot based on integration of ATAC-seq and EPIC methylation and RNA-seq data, samples represented by circles - GSD1a (red) and HC (blue), lines represent a confidence interval around each group. **e.** A table of pathway enrichment analysis results obtained from STRING clustering analysis. Clusters are shown in Figure S7d. Cluster, enriched pathway name and FDR value are shown.

### Reversal of acetylation pattern reduces the phenotypic differences between Gsd1a and HC

The deep involvement of the histone deacetylase Sirt-1 in the GSD1a phenotype and the impact of acetylation, a modification heavily dependent on metabolic status, on chromatin accessibility motivated us to investigate acetylation as a potential cause of the specific phenotype found in GSD1a patient fibroblasts. To that end, we conducted a reverse epigenetic experiment to change histone acetylation patterns in GSD1a and HC cells. We hypothesized that reversing the observed histone acetylation patterns shown above will mitigate the phenotypic differences between GSD1a and HC fibroblasts. To this end, we treated the cells with a mix of histone deacetylase inhibitors to encompass the different classes of HDACs (see Methods) that are involved in maintenance of histone deacetylations. We explored the effect of this treatment in parallel at different levels: First. We verified that the treatment did not affect cell viability by tracking the cells daily (see representative images from cell culture post treatment in Figure S10a), next, targeting H3K9 acetylation levels and downstream changes in key phenotypic markers in GSD1a cells as shown in Figure 10. Figure 10a illustrates the experimental procedure performed for this aim. Figure 10b shows the WB analysis of H3K9 levels in GSD1a and HC samples before and after HDAC inhibitors (HDACi) treatment as a way to assess the treatment’s efficacy. Before treatment the levels of acetylated H3K9 normalized to total H3 were higher in GSD1a cells as compared to HC, while HDAC inhibitors treatment significantly increased the acetylated H3K9 levels in both groups. Moreover, we assessed the downstream effect of the epigenetic HDACi treatment on key phenotypic characteristics of GSD1a fibroblasts as described above. First, we performed IF experiments measuring the intensity levels of the mitochondrial marker PGC1α and the lysosomal marker TFEB in a similar manner as described above. Our IF results suggest that the HDACi epigenetic treatment reduced the differences in these two key regulatory markers between the GSD1a and HC groups as shown in Figure 10c and Figure S10b. These results imply that changing the epigenetic histone acetylation pattern in GSD1a and HC fibroblasts samples normalized the intrinsic differences between these two groups in the expression of these two metabolic regulatory proteins. To have a more comprehensive view of the treatment effects on the GSD1a phenotype, we performed live imaging experiments as described in Figure 1 and applied a multivariate analysis. The results are shown in Figure 10d and Figure S10b. First, a heatmap of the HC-GSD1a differences in 24 selected features as selected in Figure 1. The heatmap clearly shows that differences between GSD1a and HC in these features were significantly mitigated by the reversal of histone deacetylation. The Linear Discrimination Analysis (LDA) plot (Figure 10d, bottom) shows that the treatment significantly affected GSD1a and HC cells as the treated and untreated samples were separated by LD1(86% variance explained). Interestingly, the distance between the groups in LD2 is smaller in the treated group compared to the untreated. This suggests that the HDACi treatment reduced the phenotypic differences between HC and GSD1a cells. Altogether these results show that our treatment efficiently modified the histone acetylation pattern in GSD1a and HC cells and that this treatment also decreased the phenotypic differences between these groups which strengthens our hypothesis that the phenotype in GSD1a fibroblasts is attributed to epigenetic modifications originated by an unknown mechanism in GSD1a patients.

**Figure 10.**
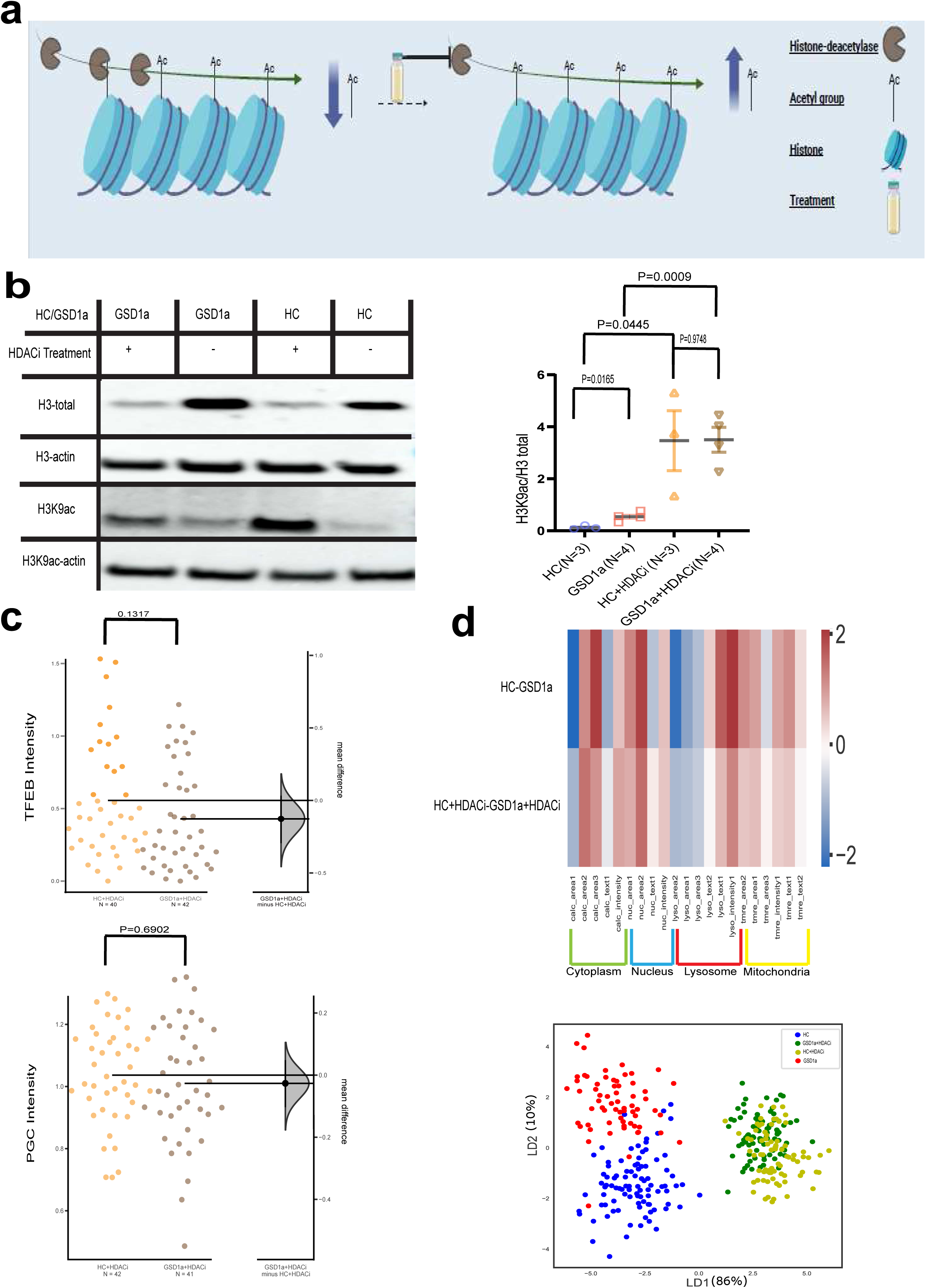
Histone deacetylation (HDACi) treatment effect on key phenotypic characteristics in GSD1a fibroblasts. **a.** Illustration of the experiment design depicting the effect of histone deacetylase inhibitors (HDACi) treatment on general chromatin histone acetylation levels. **b.** WB analysis of the effect of HDACi treatment effect on H3k9ac and H3 total levels in GSD1a and HC cells as compared to untreated samples. Shown are representative immunoblots (left) and densitometric quantifications (right). HC (blue), GSD1a (red) and HDACi treated GSD1a (brown) and HC (yellow) fibroblasts. Actin levels were used to normalize loading protein content of samples (N indicates samples analyzed for each group for ac-H3K9 divided by total H3 levels, p values were computed using two-tailed t tests). **c.** Bootstrap CI Scatter plots showing comparative analysis of TFEB and PGC1α intensity levels, following HDACi treatment, between HC (yellow) and GSD1a (brown) treated fibroblasts. The left y-axis represents protein intensity levels and the right y-axis represents mean differences between the tested groups. A 95% confidence interval derived from bootstrap resampling test is represented as a point estimate with a vertical bar and its respective distribution for the tested group effect size compared to HC (N indicates number of wells analyzed for each group, p values were computed using two-tailed t tests). **d.** Top-Heatmap analysis from image-based HCA phenotypic analysis representing 6 features for each labeled organelle. Rows presenting normalized mean difference values for HDACi untreated or treated HC and GSD1a samples in the live imaging experiment. The different feature names shown for each label are numbered for simplification. Original feature names are mentioned in Supplementary Table 1. Bottom-LDA plot, showing untreated GSD1a (Red) and HC (Blue) cells compared to HDACi treated GSD1a (green) and HC (yellow) cells.

## Discussion

Based on deep phenotypic analysis of GSD1a patients’ fibroblasts, we present here strong evidence that these cells manifest a specific disease phenotype even though G6Pase, the liver enzyme whose deficiency causes the GSD1a disease, is not expressed in these cells. This phenomenon reveals for the first time the possibility that somatic cells from a tissue which does not express a disease-causing mutated gene maintain a disease phenotype, even when kept under standard culture conditions remote from the original pathological environment. Importantly, this is a proof-of-concept work based on skin fibroblasts. Further work in other tissues both proximal and distal to the affected tissue (liver in GSD1a) will need to be performed to underpin this novel concept of disease-induced metabolites-driven epigenetic programing and systemic imprinting of a pathological status.

The effect of GHF201 treatment in GSD1a cells reverses the levels of most of the identified disease markers including glycogen, suggesting that this known glycogen-reducing agent helps improve the overall GSD1a phenotype in cells (Figure S4), while also being a systemic therapy as suggested by its ameliorating effects in a GSD1a mouse model (Figure 5). These drug effects strongly support the specificity of the GSD1a phenotype in patients’ primary fibroblasts. Moreover, the main liver features that were improved by GHF201 treatment in *L.G6pc^-/-^* mice, *i.e.,* reduction in glycogen and increase in glucose, were also replicated in fibroblasts (Figure S2a) in further support of phenocopying cell pathology in non-mutated tissues - i.e., not only cellular features, but also therapeutic responses are phenocopied. However, an important difference between liver and skin fibroblasts is that liver has a substantial accumulation and turnover of glycogen as compared to fibroblasts. Consequently, the metabolomic landscape and subsequent epigenetic programing driven by GSD1a hepatocytes are expected to be different than those driven by GSD1a skin fibroblasts. Our work only demonstrates, as a proof-of-concept, that distal tissues, such as fibroblasts, are metabolically reprogrammed by the diseased state. Our deep genotypic and phenotypic analysis demonstrate that this reprogramming is pathogenically relevant for GSD1a. However, in the organism, disease-induced metabolic modulation of other cell types, such as hepatocytes, with basal metabolism different than that of fibroblasts, would also contribute to epigenetic reprogramming and since their metabolism is different, this contribution might also be different, in terms of the specific genes modulated, than that of fibroblasts, albeit it is expected to produce similar pathological outcomes typical of GSD1a. Effectively, the epigenetic reprogramming in GSD1a, or other metabolic disorders, is a product of the disease-related metabolic perturbation of all tissues.

At this point, we have not yet identified the exact signal from the organism, nor the disease-associated programming mechanism responsible for the GSD1a fibroblasts phenotype. We are aware that this cell memory programming mechanism is different from the known genomic or behavioral imprinting concepts^52,53^. However, while this mechanism is indefinite, it is probably also multifaceted. Therefore, instead of pursuing the specific mode of action by which GSD1a fibroblasts acquire pathology, this work has three tiers: 1. Phenotype analysis of the pathology in GSD1a fibroblasts, so as to delineate the major cellular pathways it implicates; 2. Defining differentially expressed genes and metabolic pathways as putative agents of the phenotypic aberrations observed in GSD1a fibroblasts; 3. In depth analysis of putative epigenetic factors working independently and in concert to enable the environment to uniquely modify gene expression in GSD1a fibroblasts.. The main insights of our work are discussed below:

### Cell phenotyping

Image-based HCA experiments of GSD1a phenotype and deep phenotypic analysis indicate impaired lysosomal and mitochondrial functions in GSD1a cells compared to HC, which are driven by a transcriptional dysregulation of the NAD^+^/NADH-Sirt-1-TFEB regulatory axis (Figures 1,2,3,4 and S4), supported by reduced expression of lysosomal function proteins such as CTSS, CTSL (Figure 6a right) and the vATPase subunit ATP6V0D2 (Table S3). This dysregulation impacts the normal balance between mitochondrial biogenesis and mitophagy as suggested by the changes in the levels of the mitochondrial biogenesis factors PGC1α and its target genes NRF1 (Figure S3b) and PPARγ along with its coactivator subunits, alpha and beta(Table S3) and inhibition of mitophagy (Figure 4), possibly due to inhibition of the NAD^+^ and Sirt1-dependent mitophagy activators PINK1 and Parkin^54–56^ in GSD1a patients’ cells. These results correlate with the reduced expression of Sirt-1 in the livers of *L.G6pc^-/-^* mice^57^ which is a key mechanistic factor for the impaired hepatic autophagy phenotype described in this animal model, similar to the one described here in afflicted GSD1a patients fibroblasts (Figure 1 and 4). GSD1a-associated impairment of autophagy as we described here in primary fibroblasts, was also observed in hepatic cells, and in murine and canine models of GSD1a, where stimulation of the anti-autophagic mTOR and inhibition of the pro-autophagic AMPK pathways were recorded^36^. Moreover, as in our GSD1a human fibroblasts (Figure 1, 2 and S3), downregulation of *G6PC1* in *G6PC1* knockdown (KD) AML cells was associated with mitochondrial depolarization (*i.e.,* decreased TMRE fluorescence), and reduced TFAM and NRF-1 dependent mitochondrial biogenesis^58^. PGC1α levels were also reduced in livers of *L.G6pc^-/-^* knockout (KO) mice *in vivo*^58^, Remarkably, RNA-seq (Figure 6), ATAC-seq (Figure 7) and EPIC methylation (Figure 8) data, separately or integrated data (Figure 9) shows that epigenetic landmarks reflect the disease differential gene expression and metabolism expressed in the GSD1a cells which turn to be characterized as 7 different functional pathways clusters (Figure 9) that align very well with the known GSD1a pathology and metabolism (see below) and with the GSD1a fibroblasts phenotype described here for the first time.

### Key differentially expressed genes and the metabolic consequences of their differential expression

DDO is an enzyme which controls the availability of D-Asp and D-Glu that we found to be significantly decreased in GSD1a fibroblasts (Figure 6). This DDO decrease is expected to have profound implications on metabolic intracellular pathways containing D-Asp. As opposed to Asp aminotransferase (AST or GOT), which catalyzes transamination of L-Asp to L-Glu, the enantio-specific DDO catalyzes the oxidative deamination of D-Asp and partakes in peroxisomal ROS metabolism^59^. Therefore, its deficiency in GSD1a might be associated with reduced peroxisomal and ensuing mitochondrial beta oxidation as the latter is supplied by peroxisomal acylcarnitines. This indeed is in agreement with the increased levels of L-carnitine in GSD1a media (p<0.08) and with reduced expression of the positive regulator of beta oxidation PPARα. ALDH1A1 has an important role in detoxification of hepatic ethanol because it oxidizes the acetaldehyde generated by ethanol oxidation thus shifting the entire reaction towards a decrease in ethanol levels. In addition, similar to the antioxidant PPARβ, ALDH1A1 has a key role in ROS detoxification, for instance ROS produced by malondialdehyde formed by lipid peroxidation in cell membranes. Importantly, a potential damage which could be caused by ALDH1A1 deficiency is retinoic acid insufficiency, since retinoic acid (RA) is the oxidation product of retinaldehyde. This direction is especially interesting in view of the vital role of RA in early stages of embryonic development that also involve 3’ HOX gene cluster regulation resulting in expression domains that extend more anteriorly in the body^60^. RA gradients impact on gene expression during formation of multiple tissues in the embryo. A possible explanation for the failure to observe RA congenital defects in GSD1a embryos (born patients) might be because of the compensatory up-modulation of other ALDH isomers, such as the mitochondrial enzyme ALDH1L2, implicated in the folate pathway (see next paragraph), which might protect from such defects (*e.g.*, neural tube defects^61^).

A specifically interesting result is the integration between the transcriptomics and metabolomics of folate and single carbon metabolism (Figure 6d). Single carbon metabolism is the main source of S-adenosylmethionine which is the direct substrate of DNA methylation. The increase in folate (p<0.09), the significant change in one carbon pool by folate pathway, and the increased expression of ALDH1L2 as shown in Figure 6, which generate mitochondrial tetrahydrofolate, all point to substrate (folate) accumulation and down modulation of the rate of its degradation and of single carbon metabolism. Such a slowdown in single carbon metabolism might correspond to the relative global DNA hypomethylation observed in GSD1a fibroblasts (Figure 8 and S8). Trp metabolism is the only endogenous pathway for *de novo* generation of NAD^+^^62^, Trp is metabolized to kynurenine, which is then either converted to kynurenic acid in a ROS-enhanced “deadend” reaction in terms of NAD^+^ formation, or converted to quinolinic acid and then to NAD^+^^63^. The significant increase in kynurenic acid in cells, decrease in quinolinic acid in cells, and increased metabolization of the antioxidant reduced glutathione (Figure 6 and Supplementary Table 6) all suggest down-modulation of NAD^+^ formation in GSD1a fibroblasts, which is in line with the direct observation of lower NAD^+^ levels and NAD^+^-dependent Sirt-1 activity (Figure 3). These metabolites are at key nodes of the Trp metabolism pathway, which was indeed enriched in metabolite sets other than SMPDB, the KEGG pathway “Phenylalanine, tyrosine and tryptophan biosynthesis”, and the RaMP-DB pathway “Nicotinate and nicotinamide metabolism” both in cells (Figure S6c). At the RNA-seq level, decreases in IDO1, which converts indoleamine 2,3-dioxygenase 1 to Trp, PSPH, which converts quinolinic acid to NAD^+^, and the increase in kynurenine aminotransferase 3 levels (FDR<0.12), which catalyzes the deadend conversion of of kynurenine to kynurenic acid, all correspond to reduction in NAD^+^ via modulation of the Trp metabolism pathway.

### Metabolic modulation of chromatin accessibility and transcription

In line with our deep phenotyping results, here we show for the first-time strong evidence that the chromatin epigenetic signature of GSD1a cells is significantly different than that of HC cells. This signature can be demonstrated by the differences in open chromatin regions (Figure 7 and S7), by the different global patterns of non-methylated regions of genomic DNA (Figure S8a), by pattern differences in major methylation sites (Figures 8 and S8) and by high histone 3 acetylation levels in different sites (Figures 3, 10) which may indicate global dysregulation of histone acetylation that have a direct downstream effect on gene expression and cell phenotype as discussed below.

A key question is what the source of these epigenetic modifications is. It is generally accepted that epigenetic changes of both DNA and histones are mostly generated during embryonic development by the local cell environment^64^. However, epigenetic reprogramming is also a part of embryonic differentiation. Following fertilization, both maternal and paternal DNAs are passively demethylated during cell replications, while paternal DNA is also actively demethylated by TET proteins^65^. *De novo* methylation and methylation maintenance, on the other hand, are mediated by methyltransferases from the blastocyst stage throughout embryonic development. Both methyltransferases (‘writers’) and demethylases (‘erasers’) are tightly regulated. The dependence of methyltransferases on acetylation^66,67^ and of the TET demethylases on N-acetylglucosamine, phosphorylation, acetylation, and α-ketoglutarate^68,69^ suggests that this regulation depends on extracellular metabolites.

Due to the deamination of methylated cytosine to thymine and the methylation-dependent silencing of transposable elements^70^, DNA-methylation is significantly mutagenic^71^. In addition, loss-of-function mutations in many genes are associated with increased methylation of promoter and enhancer sequences of these genes. This phenomenon has especially been described in cancer, where hypermethylation of promoters of tumor suppressor genes is a known tumorigenic factor. Promoter hypermethylation is also demonstrated, for instance, in downregulated genes in the plant *F. fusca*^72^, in inhibited lung pro-inflammatory genes^73^, and in loss of function of the fragile X mental retardation 1 (FMR1) gene^74^. Interestingly, our own data also shows hypermethylation at the genomic site of the *G6PC1* (Figure S8d). Regardless of this mechanism, we found some evidence for an epigenetic change in GSD1a fibroblasts. This change is in classical developmental HOXC genes in chromosome 12. Several differentially methylated regions (DMR) are found in proximity to to the region where concomitant down-regulation of HOXC gene cluster expression might occur, and where strong differences in the methylation pattern between GSD1a and HC samples are clearly visible across the chromosome’s regions (Figure 8c and Supplementary Table 9). Functionally, in contrast to HOXC cluster heavy down regulation we found the HOXD homeobox cluster genes in Chromosome 2 to be up-regulated (Figure 6b). Moreover, a broad set of regulatory RNA genes - HOXC-AS1, HoxC-AS3, HoxC-AS2, HoxC13-AS, HoxB-AS3 and HOTAIR that are affiliated with the lncRNA class were downregulated as well (Figure 6b), strongly supporting broad silencing of HOXC and HOXB families. These type of HOX clusters gene repression may function in a similar way as known for HOTAIR, which binds lysine specific demethylase 1 (LSD1) and polycomb repressive complex 2 (PRC2) and serves as a scaffold to assemble these regulators at the HOXD gene cluster, thereby promoting epigenetic repression of HOXD^75^.

### Metabolic-driven, disease-associated programming of cell memory

Beyond the above-mentioned general mechanism regulating methylation and acetylation, our results also suggest a GSD1a-disease-specific metabolic mechanism for programming cell memory in all tissues expressing the G6PC1 mutant with excess glycogen: glycogen binds the ß-subunit of the anti-cancer protein AMPK and inhibits it. In GSD1a cells which do not express the G6PC1 mutant, such as fibroblasts, p-AMPK can be inhibited by the reduction in NAD^+^ (Figure 3b) with ensuing inhibitions of Sirt1 and the direct AMPK kinase LKB1. As an essential component of the p-AMPK-Sirt-1-NAD^+^/NADH axis, p-AMPK drives histone deacetylation (*e.g.* ^74^) a key epigenetic characteristic which was inhibited in GSD1a cells (compared to HC) by inhibition of the AMPK-Sirt-1 axis (Figure 3). These observations suggest that the pathological accumulation of glycogen in GSD1a patients drives the epigenetic programing of the disease memory in the pathogenic, glycogen accumulating tissues (liver and kidney), while epigenetic programming is mediated by other metabolic or redox pathways, such as NAD^+^/NADH modulation. Thus, in GSD1a the diseased state, manifested as a unique intracellular metabolic aberration, *i.e.* over-accumulation of glycogen, or compromised OCR and therefore lower NAD^+^/NADH, could program the memory of the disease in all cells of the organism through the p-AMPK-Sirt1-NAD^+^/NADH axis. This programing is probably mediated by different mechnisms in cellsexpressing or not the G6PC1 mutant. We term this new phenomenon *Disease-Associated Programming,* and we conjecture that this is a novel mechanism by which the memory of a cellular metabolic aberration can be programmed in all different organismal tissues and throughout embryonic development.

In support of this concept, evidence from studies investigating the link between metabolism and epigenetic changes in disease show similar alteration in relevant energy metabolites that play as cofactors for chromatin modification of epigenetic markers such as dysregulation of histone acetylation and DNA methylation patterns as described in our work^76^. An interesting example from the work of Katarzyna et al., showed the direct effect of a ketone metabolite on cellular bioenergetics which in turn elevates acetylated histone 3 levels required for CD8^+^ T cell activation^77^. Another example of a connection between metabolism and epigenetics was shown in HepG2 cells were the expression of gluconeogenic and glycogenolytic genes was affected by decreased levels of the histone modifying enzyme LSD1 (Lysine-specific demethylase)^78^. Moreover, blood samples of individuals (∼700) suffering from metabolic syndrome (elevated fasting plasma glucose level and increased cholesterol and triglycerides) showed global reduced DNA methylation supporting the DNA methylation profile found in GSD1a patients’ fibroblasts which also can suffer from similar plasma metabolic dysregulation^79^. Interestingly, hypomethylation also supports activation of pro-glycolytic enzymes required for aerobic glycolysis (Warburg effect) in cancer^80–82^ evoking investigation of the link between metabolically induced hypomethylation, induced glycolysis (Figure 2i) and tumorigenesis or incidences of adenocarcinoma in GSD1a^83^. This link is particularly plausible since highly pro-glycolytic genes such as CXCR4^84^ and CTHRC1^85^ were found to be significantly hypomethylated in GSD1a patients’ fibroblasts (Supplementary table 8). In addition, in depth studies on epigenetics in human obesity and type 2 diabetes revealed a distinct DNA methylation pattern based on the Infinium 450K array as we found in our study^86^. In terms of embryonic development, our newly proposed *Disease-Associated Programming* posits that a mutation, such as *G6PC1^-/-^* is initially expressed at the blastocyst stage^19,20^, in the same way that all genes are expressed in a non-regulated manner. This initial expression of the metabolic mutant would irreversibly imprint aberrant epigenetic changes, which would be carried over throughout development, overriding other developmental differential expression mechanisms, which might also be epigenetic or environmental, such as the differential suppression of the mutant expression in certain cell lineages.

### Differential acetylation as a unique driver of disease-associated programming

To explore the hierarchical relationship between the epigenetic and the cellular phenotypic characteristics mentioned above, we performed a broad deacetylation inhibitory experiment aimed to induce similar acetylation patterns in HC and GSD1a cells while not affecting their cellular viability (Figure 10 and Figure S10a and b). We show in these experiments that this treatment significantly reduced the disease phenotype found in GSD1a cells as compared to treated HC. This result strengthens our hypothesis that GSD1a cells have a metabolically-regulated epigenetic phenotype which affects downstream multiple cellular functions and pathways. Interestingly, this HDACi treatment effect also indicates the epigenetic reversibility of the phenotype which can be supported by a recent seminal work^87^. This work presents a novel transient naive reprogramming treatment of primary skin fibroblasts which can reset epigenetic reprogramming to the pre-implantation stage, bypassing the erasement of epigenetic imprinting which occurs at the post-implantation stage and is common with erasement of this imprinting or memory by the Yamanaka method used to generate iPSC^88^ of somatic cells. We estimate that the more genuine transient naive treatment (TNT) epigenetic reprogramming method, recapitulating the natural human embryonic stem cells (hESC), rather than human induced pluripotent stem cells (hiPSC), can better preserve imprinting memory.

### Metabolism as a driver of epigenomic landscape: G6PC1 non-expressing GSD1a fibroblasts revive a historic model

A highly relevant review on the impact of cellular metabolism on chromatin dynamics and epigenetics mentioned the *avant-garde* work of Waddington in the 1940’s, describing for the first time the concept of epigenomic landscape to illustrate the relationship between the epigenome and different cell states^89^. This landscape is composed of valleys and summits, the valleys represent the different cell phenotypes, which, in our case, are GSD1a and HC fibroblasts, while the summits correspond to the barriers that maintain the differences between epigenetic phenotypes. Waddington’s epigenomic landscape concept proposed two different metabolic reprogramming models that facilitate the transition between epigenomic landscapes. One model claim that metabolism facilitates cell state transitions while the other claims formation of new cell types through the same metabolic changes. Applying our results into this perspective, we think that the metabolic landscape originated by the G6PC1 mutation in liver and kidney cells in GSD1a patients has led to metabolic reprogramming (probably during embryonic development) changing the epigenomic landscape in GSD1a fibroblasts which is distinct from that of HC fibroblasts. Interestingly, another recently published paper unveils a similar message of metabolically-driven epigenetic reprogramming: In their recently published work show that intermittent fasting generates an epigenetic footprint in PPARα-binding enhancers that is “remembered” by hepatocytes leading to stronger transcriptional response to imposed fasting by up-regulation of ketogenic pathways. In the same way, the diseased GSD1a status in our case imposes metabolic changes, as detailed here, leading to permanent epigenetic changes, also described here, which are “remembered” by GSD1a fibroblasts and which play a major role in the transcription of pathogenic genes, which is how the diseased state is preserved even in cells not expressing the G6Pase mutant, which is the direct cause of the disease.

### Summary and outlook

The meaning and importance of a cell memory mechanism in somatic cells like GSD1a skin fibroblasts that allegedly do not comprise the tissue affected by the disease is that these cells “remember” the diseased state and respond to external conditions and therapeutic cues in the same way as primary affected cells, such as liver cells. This behavior implies that cells in the entire organism are affected by the disease and not only the cells expressing the mutated gene, Intriguingly, this new concept might lead to a full revision of thoughts on the efficacy and future of tissue-targeted gene therapies like those being developed for GSD1a^91,92^. In extrapolation, we anticipate that, being based solely on the Sanger sequence of affected genes, gene therapy will not be able to erase the disease signature from fibroblasts or other tissues which are not genetically implicated in the disease, but which might contribute to its pathology by their disease-associated programming. For GSD1a, it is probable that potential GSD-curing small molecules such as GHF201 could complement putative gene therapy by supplementing the genetic treatment with an active glycogen or NAD^+^ reducing effect to reverse the established pathology in pathogenic and peripheral patients’ tissues.

## Acknowledgments

We would like to thank the following people: Dr. Yuval Ebenstein and Dr. Rachel Zamostiano, Tel Aviv University for their advice and technical support; Yuval Raviv for the initial contribution to the project; Roi Ronen for the comments and advice on the algorithms used; and Dr. Ido Goldstein from the Hebrew University of Jerusalem for the Histone H3K27ac antibody. The PhD scholarship of US was partly supported by The Yoran Institute of Genome Research and Personalized Medicine and by The Tel Aviv University Prajs-Drimmer Institute for Development of Anti-Degenerative Drugs. This work was funded by the Krigsner Foundation and is dedicated to the memory of David Krigsner Zindeluk (z”l).

## Author contributions

SU, DSJ, MK, MN designed and performed most experiments; CMA and MG designed and performed the in vivo experiments; CF and KH performed and participated in the DNA ATAC seq and genomic integration data analysis, respectively; AS performed and participated in the data analysis of SIGL; SU supervised and performed the data analyses throughout the article; MG, RF, BY and LM supervised experiments and critically reviewed the manuscript; AY contributed with clinical samples and reviewed the manuscript; KO and WM coordinated and supervised all experiments; SU, KO and WM wrote and reviewed the manuscript.

## Methods

### Chemicals and reagents were purchased from Merck-Sigma-Aldrich or as otherwise stated

#### Patients and cell culture

Primary Skin fibroblasts were isolated from skin biopsies of 5 GSD1a patients (age 5-25, See Table 1). The patients were on a dietary treatment which includes nocturnal nasogastric infusion of a high glucose formula in addition to usual frequent meals during as advised in ^93^, Patient 6894 did not tolerate well the uncooked cornstarch and therefore was treated with a tailored dietary treatment planned by metabolic disease specialists and dedicated certified dieticians highly experienced with the management of pediatric and adult patients with GSDs and other inborn errors of metabolism. The biopsies of patients were taken within the range of 3 month to several years upon receiving the aforementioned dietary regimen. Informed written consent to participate in the study was obtained from all individuals or their parents in the case of minors. The study was approved by the ethics committee of the Institutional Review Board of the Sheba Medical Center (5722-18-SMS). Age and gender matched healthy controls (HC) were purchased from Coriell Institute (New Jersey, USA) (See Table 1 below). Primary fibroblasts were then expanded in culture media DMEM supplemented with 1% MEM Sodium Pyruvate, 1% PSA (Biological industries, Israel), 10% heat inactivated FBS, and 1% 100x non-essential amino acids solution (NEAA) in polystyrene plastic 75-cm2 culture flasks (Corning, NY) at 37°C with 5% CO2. Cells passages and expansion of human skin fibroblasts were performed when cells were at 80-100% confluency. Cells passaging and re-plating was accomplished using 0.25% Trypsin-EDTA (Biological industries, Israel) for 2 minutes, followed by addition of twice the volume of a complete culture media to neutralize the enzyme. Cells were subsequently centrifuged at 1200 rpm for 5 minutes before their pellets were resuspended in 1 ml full medium for cell counting using TC10 automated cell counter (BioRad USA) before re-plating. All GSD1a and HC samples (See Table 1) were used for each experiment unless mentioned otherwise. All experiments were performed between passages 8– 16 as this range was determined as an adequate passage range in previous studies^13,14^.

**Table 1.** Cells.

| Cell Type | Cell id | Age | Gender | Mutation (G6PC1 gene) |
| --- | --- | --- | --- | --- |
| GSD1A | 0762 | 5 | Male | c.247C>T, p.Arg83Cys and<br>c.827G>T, p.(G276V)( *SI gene* ) |
| GSD1A | 75858 | 19 | Male | c.247C>T, p.Arg83Cys |
| GSD1A | 361 | 15 | Female | c.247C>T, p.Arg83Cys |
| GSD1A | 6894 | 25 | Male | c.247C>T, p.Arg83Cys |
| GSD1A | 63852 | 14 | Female | c.247C>T, p.Arg83Cys |
| HC | GM00498 | 3 | Male |  |
| HC | GM00038 | 9 | Female |  |
| HC | GM07522 | 19 | Female |  |
| HC | GM00499 | 8 | Male |  |
| HC | GM09503 | 10 | Male |  |
| HC | GM17507 | 16 | Male |  |
| HC | GM03377 | 19 | Male |  |
| HC | GM05400 | 6 YR | Male |  |

#### Live Imaging

GSD1a and HC skin fibroblasts were seeded at 1,400 cells per well and cultured in specialized microscopy□grade 96□well plates (Cellvis <u>P96-1.5H-N</u>). Following 24 h in complete medium (DMEM with glucose,10% FBS, PSA, NEAA and sodium pyruvate), cells were washed to remove traces of serum and starvation medium (DMEM serum free, glucose and pyruvate free, with PSA) was applied for 24 h and 48 h or for 48 h followed by 24 h with complete medium (72 h). Before imaging, plates were examined for well quality using a light microscope, following that the cells at the end of their respective time points were washed and a mix of live fluorescent dyes in HBSS was added to each well for 20 min at 37°C in a 5% CO_2_ incubator. The final solutions of the different stains were: 1.6mM Hoechst 33342, 0.1 mM LysoTracker Deep Red, 0.05 mM TMRE and 0.4 mM calcein-AM Green (Thermo-Fisher Scientific). Following the incubation, cells were washed with HBSS, and plates were transferred to an Operetta G1 system for image acquisition at 20× magnification under environmental control conditions at 37°C and 5% CO_2_. All the assay parameters (including the acquisition exposure times, objective, and the analysis parameters) were kept constant for all assay repetitions. These experiments were repeated for three consecutive weeks, for a total of 18 plates (6 per condition), each plate containing 60 wells with ∼500 cells per well. Images were analyzed by a customized image analysis protocol designed for the experiment using Harmony 4.8 image analysis software, all data analysis steps were performed using Python and R. Image analysis and downstream analysis was guided by Caicedo et al^94^ and included image processing, organelle segmentation, quality control assessment for each plate, batch effect analysis, well removal based on extreme values, removal of outlier wells, feature selection by backward feature elimination, logarithmic normalization to achieve gaussian distribution levels prior to statistical analysis and multivariate analysis.(Figure S1a, Data Analysis and statistics below)

#### Mitochondrial activity assays

Mitochondrial respiration (OCR and ECAR) of primary skin fibroblasts was quantified in real-time using the Seahorse extracellular XFe96 flux analyzer (Agilent). Cells were plated in XFe96 cell culture plates at a density of (40000 cells/well), cells were incubated for 48 h in starvation medium, followed by the replacements with complete medium for 24 h as described above, 50 µM GHF201 treatment was added to half of the GSD1a wells. Cells were then washed in assay media (XF Base media (Agilent) with glucose (10 mM), sodium pyruvate (1 mM) and L-glutamine (2 mM) (Gibco), pH 7.4 at 37 °C). OCR and ECAR were measured using Seahorse’s Mito Stress assay (Agilent), with the preset automatic addition of oligomycin (1.5µM), carbonyl cyanide 4-(trifluoromethoxy) phenylhydrazone (FCCP; 2 µM) and antimycin A and rotenone (0.5 μM each)). For ATP fluxes from glycolysis and oxidative phosphorylation estimation in the cells, Agilent’s Seahorse Real-Time ATP assay kit was used. Cells were seeded and serum/glucose-starved for 48 h and the full medium was then replenished for 24 h without (UT) or with (chronic treatment) 50 µM GHF201. For acute treatment, 50 µM GHF201 was added to the for 20 min after 24 h of serum/glucose replenishment. The Real-Time ATP Assay employs a sequential injection of Oligomycin and Rotenone/Antimycin A. Agilent’s Seahorse XF Glycolysis Stress Test was performed using an XFe96 analyzer. Cells were treated with 50 µM GHF201. The assay consists of the subsequent addition of glucose, oligomycin and 2-DG and measurement of the key parameters of the glycolytic function as determined by the manufacturer’s protocol.

#### Immunofluorescence

HC and GSD1a cells were cultured, 1800 cells per well were seeded in specialized microscopy□grade 96 well plates (Cellvis <u>P96-1.5H-N</u>) and cultured overnight for attachment. Cells were incubated for 48 h in starvation medium, followed by the replacements with complete medium for 24 h as described above, 50 µM GHF201 treatment was added to half of the GSD1a wells. Cells were washed in PBS and then fixed in 4% paraformaldehyde in PBS, washed three times in PBS, permeabilized in 0.1% Triton X-100 in PBS for 5 min and blocked-in blocking buffer (5% FBS & 2% BSA in PBS) for 1 h. Cells were then incubated with primary antibodies (See Table 2 below) overnight, followed by three washes in 0.05% Triton X-100 in PBS and incubation with secondary antibodies and counterstaining with 1.6mM Hoechst 33342 (1:1000) and 0.02 mM phalloidin Atto 565 (1:400) for 1 h. Cells were subsequently washed three times in PBS. Images were acquired with 20x magnification using the Operetta High content imaging system. The experiment was repeated 3 times per primary antibody where the assay parameters (including the acquisition exposure times, objective, and the analysis parameters) were kept constant for each antibody. Images were analyzed by the Harmony 4.8 image analysis software using a customized image analysis protocol and quantified for intensity levels of each antibody. Each plate was normalized using the maximal intensity levels to achieve normally distributed features for statistical testing, and then the similar plates were combined and normalized to HC levels to perform statistical analysis. Statistical analysis for the p-AMPK/AMPK experiment was performed on the ratio of p-AMPK to total AMPK in each sample tested. Signal Outlier value detection was used to remove outlier wells using the Prism Route method (Q=10%).

**Table 2.** Antibodies.

| Antibody | Company | Dilution | Method |
| --- | --- | --- | --- |
| Rb- NRF1 (ab34682) | Abcam | 1:1000 | WB |
| Rb-TFAM (D5C8) | Cell Signaling<br>Technology | 1:1000 | WB |
| Rb- COXIV (3E11) | Cell Signaling<br>Technology | 1:1000 | WB |
| Goat-PGC1α (ab106814) | Abcam | 1:50 | IF |
| Rb-TFEB(D207D) | Cell Signaling<br>Technology | 1:100 | IF |
| Rb-P62(109012) | Abcam | 1:150 | IF |
| Rb-LC3(2775S) | Cell Signaling<br>Technology | 1:1000 | WB |
| Rb- Sirt-1(ab189494) | Abcam | 1:100 | IF |
| Donkey anti-rabbit Alexa Fluor 488 conjugated | BETHYL | 1:1000 | IF |
| Donkey anti-goat Alexa Fluor 488 conjugated | Thermo | 1:1000 | IF |
| HRP-conjugated donkey anti-rabbit | Abcam | 1:10000 | WB |
| Histone H3K27ac antibody (pAb) Rabbit anti immunogenic peptide | Active Motif | 1:200 | IF |
| Histone H3K9ac -Acetyl-Histone H3K9 Rb monoclonal antibody (mAb) (9649T) | Cell Signaling Technology | 1:1000 | WB |
| Histone H3 Rb monoclonal antibody (mAb) (4499T) | Cell Signaling Technology | 1:1000 | WB |

#### Lysosomal pH ratiometric measuring

Ratiometric analysis of the Lysosensor dye staining for lysosomal pH was performed by ratiometric fluorimetry using the BD-LSRII flow cytometer as described previously^13^.

#### Immunoblotting

HC and GSD1a cells were cultured, seeded in 6 well culture plates and cultured overnight for attachment. Cells were incubated for 48 h in starvation medium, followed by the replacements with complete medium for 24 h as described above, 50 µM GHF201 treatment was added to half of the GSD1a wells. Cells were washed in PBS and harvested and lysed in a RIPA buffer containing phosphatase and protease inhibitor cocktails. Protein concentration was determined using a Pierce BCA protein assay kit (Thermo Fisher Scientific). Briefly, an equal amount of protein lysates was subjected to SDS-PAGE 4– 12% gradient gels and electro transferred to a nitrocellulose membrane (iBlot 2, Invitrogen). Membranes were blocked with 5% non-fat milk, incubated at 4 °C overnight with primary antibodies (See Table 2 below), washed, and then incubated with secondary antibodies conjugated to HRP. Proteins were then visualized by chemiluminescence reagent (Cyanagen (XLS3,0100), Thermo Fisher Scientific (34580)) using Amersham Imager 600 (GE Healthcare) and band densitometry was measured using the FIJI software. Experiments were repeated independently three times. Protein bands were normalized to Actin levels (See Table 2 below) in each membrane.

#### Autophagy Flux calculation

Autophagy Flux was calculated by the formula below using the data acquired from different blots for each sample averaged across experiment replicates as shown in Figure 4. 50uM Vinblastine (Cayman,11762) was applied 2 hours prior to protein extraction.

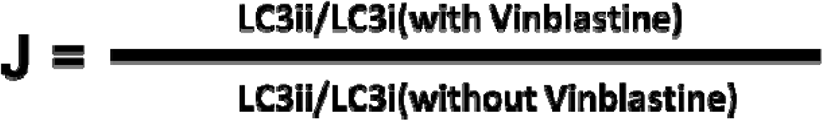

#### Periodic Acid Shiff (PAS) assay

For PAS staining of glycogen, GSD1a skin fibroblasts were seeded at 1,400 cells per well and cultured in microscopy□grade 96□well plates (Cellvis <u>P96-1.5H-N</u>) for 24 h in complete medium, 50 µM GHF201 treatment was added to half of the GSD1a wells.Cells were washed with PBS and then fixed and permeabilized with 0.1% Triton X□100 as described above. PAS staining was performed using the Merck PAS assay kit (cat.101646), in a similar way but without amylase treatment as previously described by us^12^. The PAS-stained cells were imaged using InCell2200 (GE Healthcare, U.K.) with Texas red channel filters. Image analysis for cellular PAS staining was performed using the In Carta image analysis software. Total intensity was measured, normalized to HC levels, and compared between groups. The experiment was repeated independently three times.

#### NAD^+^/NADH measurements

GSD1a and HC skin fibroblasts were seeded at a density of 10,000/well in a 384 white-walled tissue culture plate (Greiner bio-one-E21123EA). Cells were cultured overnight for attachment, then cells were incubated for 48 h in starvation medium, followed by the replacements with complete medium for 24 h as described above, 50 µM GHF201 treatment was added to half of the GSD1a wells. Cells were washed in PBS and the final volume per well was 50ul. On the day of the experiment the plate was equilibrated at room temperature for 5 minutes. 50ul of NAD^+^/NADH-Glo™ Detection Reagent (Promega) was added to each well. Plates were gently and briefly shaken to mix and lyse the cells. The cells were incubated with the NAD^+^/NADH-Glo™ Detection Reagent for 30–60 minutes at room temperature. The luminescence was recorded using a Berthold Mithras luminometer. In parallel, the same batch of cells was also subjected to dynamic Incucyte analysis of cell confluency and cell death over time.

#### Global unmodified CpG pattern analysis-Simultaneous global labeling (SiGL)

Global unmodified CpG islands were characterized using the methodology described previously^95–97^.

#### DNA construct and expression for the mitophagy assay

A plasmid encoding mCherry, GFP and residues 101 to 152 of human FIS1 (mCherry-GFP-FIS1 101 to 152) was obtained from University of Dundee (MRC-PPU reagents, DU40799)^28^. cDNA for mCherry, GFP and residues 101–152 of human FIS1 were cloned into a pBABE.hygro vector. The construct was co-transfected into 293FT cells with GAG/POL gpt and VSV-G expression plasmids. For retrovirus production, calcium phosphate precipitation method was used. Supernatant containing retrovirus was collected 48 h after transfection and was filtered using a 0.45 mm filter. HC and GSD1a fibroblast cells were infected with filtered viral supernatant containing 8 mg/ml polybrene. 2 days after the infection the cells were allowed to grow, and the experiment was performed as follows:

#### Mitophagy assay

Cells stably expressing mCherry-GFP-FIS1-leading sequence^98^, were plated on a microscopylJgrade 96 well plate and incubated for 24 h in complete medium, 50 µM GHF201 treatment was added to half of the GSD1a wells. The next day, cells were treated with 4 µM FCCP for 20 minutes. Cells were then washed with PBS, fixed with 4% paraformaldehyde, washed thrice, incubated for 10 min with 1.6mM Hoechst 33342 (1:1000), and washed thrice with PBS. Plates were then transferred to an Operetta G1 system for image acquisition at 20× magnification in 37°C and 5% CO_2_. Images were analyzed using the Harmony 4.8 image analysis software using a customized image analysis protocol. Each plate was normalized using the maximal intensity levels. Then similar plates were combined and normalized to HC levels to perform statistical analysis. Statistical analysis was performed on the ratio of mCherry to mCherry+GFP signal. Outlier value detection was used to remove outlier wells using the Prism Route method (Q=10%).

#### *In vivo* experiments

Liver *G6PC1* KO was induced in L-G6PC^-/-^ mice using a CRE recombinase fused to a mutated ligand-binding domain of the estrogen receptor, activated by tamoxifen treatment, as described in^29^. Briefly, a group of 5 male mice per arm (6–7-week-old) were treated by 0.2 mg/day of tamoxifen. One month after *G6PC1* deletion, ND-GHF-201 was administered daily *per os* at 70 mg/mL. After 4 weeks, mice were starved for 6 h, euthanized by cervical dislocation and liver and blood samples were collected for further analyses. Liver glucose-s-phosphate, glycogen and glucose and blood glucose were determined as previously described^29^. Mice were housed in the animal facility of Lyon 1 University (Animaleries Lyon Est Conventionnelle et SPF) in controlled temperature (22 °C) conditions, with a 12-h light-12-h dark cycle and free access to water and standard rodent chow. Fasting was conducted with free access to water. All procedures were performed in accordance with the principles and guidelines established by the European Convention for the protection of Laboratory Animals and were approved by the local animal ethics committee and the French Ministry of National Education, Higher Education and Research (Apafis Permit #30629).

#### RNA-seq

3 GSD1a and 3 HC cells were grown in 75 cm flasks. Following 100% confluency of the cells, each flask was supplemented with a complete medium at 37°C with 5% CO_2_ overnight.Then, cells were treated for 48 h in starvation medium, followed by the replacements with complete medium for 24 h as described above. Following that cells were extracted from each flask and RNA extraction was performed using TripleXtractor Direct RNA Kit (Cat# GK23.0100) according to the manufacturer instructions. Library preparation was performed using xGEN Broad-range RNA Library Prep (IDT, Cat# 100009813) in accordance to the manufacturer instructions, Poly A mRNA Magnetic Isolation Module (Cat# E7490L) was performed under these sequencing conditions -FCL PE100 (Read length: 2×100) using a flow cell of 1500M reads*2. Sequencing was performed using DNA SEQ-G400 from MGI pair-ended. RNA extraction, library preparation and sequencing were performed by IDT (Israel). Fastq files were generated and quality control of files was performed using fastqc(0.12) and multiqc softwares^99,100^. Fastq files were trimmed using Trimmomatic (0.39)^101^ followed by another fastqc and multiqc report. Alignment was performed to the GRCh38 human genome using STAR (2.7.11b)^102,103^. Technical biological replicates were merged and the output files were sorted, indexed and used for calculation of mapping statistics using SamTools(1.18)^104^. PCR-duplicates were removed using Picard MarkDuplicated (4.0.1.1)^105^. Gene counts were generated using HTSeq 2.0^106^. Further processing of counts data was performed using R. First, QC was performed by measuring counts statistics and visualizing sample count distributions using boxplots and histograms (See Figure S5a). Counts were normalized using DESeq2^107^ and filtered for genes with counts with a mean of >40 across samples to reduce artifacts and low count genes. Annotation for genes was performed using AnnotationDbi^108^.

#### Metabolomics

3 GSD1a and 3 HC cells were grown in 75 cm flasks. Following 100% confluency of the cells, each flask was supplemented with a complete medium at 37°C with 5% CO_2_ overnight. Then, cells were treated for 48 h in starvation medium, followed by the replacements with complete medium for 24 h as described above. Subsequently, media was extracted from each flask. Following that, cells were extracted from each flask and frozen in -20C. LCMS metabolomics analysis was performed as follows. Dionex Ultimate 300 high-performance liquid chromatography (UPLC) system coupled to an Orbitrap Q-Exactive *plus* Mass spectrometer (Thermo Fisher Scientific) with a resolution of 70,000 at 200 mass/charge ratio (m/z), electrospray ionization in the HESI source, and polarity switching mode to enable both positive and negative ions across a mass range of 70 to 1000 m/z, was used. The UPLC setup included a ZIC-pHILIC column (SeQuant; 150 mm × 2.1 mm, 5 μm;Merck) with a Sure-Guard filter (SS frit 0.5 μm). Five µL of the cell extracts were injected and the compounds were separated with a mobile phase gradient of 15 min, starting at 20% aqueous (20 mM ammonium carbonate adjusted to pH 9.2 with 0.1% of 25% ammonium hydroxide) and 80% organic (acetonitrile) and terminated with 20% acetonitrile. The flow rate and column temperature were maintained at 0.2 mL/min and 45 °C, respectively, for a total run time of 26 min. All metabolites were detected using mass accuracy below 5 ppm. Thermo Xcalibur was used for the data acquisition. Data processing and analysis were carried out using Compound Discoverer version 3.3.2.31. in total, 183 metabolites were identified through this platform. The identification process utilized an in-house spectral library and MS2 data comparison with the mzCloud database. Metabolomics data was analyzed using Metaboanalyst 5.0^109^. Metabolites with more than 33% missing values were removed, metabolites that were near-constant throughout the groups were detected using a 20 IQR variance filter. Samples were normalized to the HC group and scaled using logarithmic normalization (See Figure S6a).

#### Global epigenetic modification of histone deacetylase activity

4 GSD1a and 4 HC cells were grown in two 75 cm flasks. Following 100% confluency of the cells, each flask was split into 2 different 75 cm flasks supplemented with complete medium at 37°C with 5% CO_2_ overnight. Then, cells were treated for 72 h in complete medium containing an epigenetic modifying cocktail mix of histone deacetylases inhibitors-0.5uM Vorinostat, 0.5uM Sirtinol. Medium and treatment were replaced every 24 h to ensure the effectiveness of the treatment on the cells. In addition, cell viability was measured daily (see Figure S10a). After treatment, one flask from each sample was used for protein extraction that was used for histone acetylation analysis and one flask was used for live imaging and IF analysis in a similar way as explained above.

#### Methylation EPIC array

4 GSD1a and 4 HC cells were grown in 75 cm flasks. Following 100% confluency of the cells, each flask was supplemented with a complete medium at 37°C with 5% CO_2_ overnight. Then, cells were treated for 48 h in starvation medium, followed by the replacements with complete medium for 24 h as described above. Subsequently, DNA was extracted from each flask using Qiagen’s DNeasy Blood & Tissue Kit (Cat. No. / ID: 69504), quantified, and assessed by nanodrop and sent to for the execution of the EPIC array experiment at Yale University (Yale Center for Genome Analysis). Output results were processed and analyzed using R and genome studio. Quality control and processing steps were performed initially using Illumina’s Genome studio software (see Figure S10c and Supplementary Table 12). Further quality control steps included detection of a p-value for every CpG in every sample, which is indicative of the quality of the signal for each sample, filtering out probes from the X and Y chromosomes of each sample, removal of all probes where common SNPs may affect the CpG and removal of poor performing probes based their detection p-value. Normalization method was chosen following implementation of the Quantro package analysis protocol^110^. Normalization of the data was performed using Functional normalization (FunNorm) which removes unwanted variation by regressing out variability explained by the control probes present on the array. Following that M and B values were generated for statistical analysis^110–114^. (See Figure S7c)

#### Assay for Transposase-Accessible Chromatin using sequencing (ATAC-seq)

2 GSD1a and 2 HC cells were grown in 75 cm flasks. Following 100% confluency of the cells, each flask was supplemented with a complete medium at 37°C with 5% CO_2_ overnight. Then, cells were treated for 48 h in starvation medium, followed by the replacements with complete medium for 24 h as described above. ATAC-seq library preparation was performed on 2 HC and 2 GSD1a cells using the ATAC-seq kit from Active motif (#13150). Cells were frozen with freezing media (FBS with 10% DMSO). After thawing, cell viability was assessed using Trypan Blue 0.5% solution (1:1) (Sartorius 03-102-1B) and automated cell counter (Bio Rad TC20). 100,000 live cells from samples with cell viability above 85% were used for library preparation performed according to the manufacturer’s recommendation. After the library preparation, quality control was performed to evaluate the size distribution via High Sensitivity D1000 ScreenTape and Tapestation Analysis Software 4.1.1 (Agilent Technologies). DNA quantification was performed using Qubit dsDNA HS Assay kit (Thermo Fisher Scientific). Sequencing was performed using NovaSeq 6000 v1.5, 2 x 111 pair ended. Fastq files were generated and quality control of files was performed using fastqc(0.12) and multiqc softwares^99,100^. Illumina adaptor sequences were trimmed using Trimmomatic (0.39)^101^ followed by another fastqc and multiqc report. Technical biological replicates were merged. Reads were then aligned to the GRCh38 human genome using Bowtie2 (2.5.1)^102,115^. Output files were sorted, indexed and used for calculation of mapping statistics using SamTools(1.18)^104^. Reads mapping to mitochondrial-dna were removed with SAMtools(1.18)^104^. PCR-duplicates were removed using Picard MarkDuplicated (4.0.1.1)^105^. Files were then visualized using Integrative Genomics Viewer (IGV) (2.4.9)^116^, the peaks from individual biological replicates were called with MACS2(2.2.9.1)^117^. Further processing of the peak data, QC, normalization and counting of peaks was performed using the R package DiffBind (3.17)^118,119^. Peaks were normalized based on peak library size and then annotated to the human genome using the R package Chipseeker^120,121^. Normalized BigWigs from merged files were generated for visualization using DeepTools (3.1.3)^122^, which were merged for visualization via Integrative Genomics Viewer. Visualization of the ATAC-seq QC is shown in Figure S6c. Motif analysis was performed on up/down regulated peaks (FDR<0.1) using HOMER^123^ using hg38 genome and size -200.

#### Statistics

Generally, in all experiments the number of primary fibroblasts samples used varied between 3 – 5 for the GSD1a group and 3-6 for the HC group. For calculating p-values, statistical testing by 2 tailed unpaired t-tests was performed to compare groups using GraphPad Prism 9, unless otherwise mentioned. Throughout the manuscript p values under 0.1 were considered significant, we present the actual p values for full transparency and as we strongly believe p values require context and not a defined threshold. In addition, as extremely small sample sizes such as shown in this manuscript result in lower statistical power, a higher *p*-value threshold allowed us to identify potentially meaningful differences between groups. In addition, we have focused most of the statistical analysis in this manuscript on 95% bootstrap mediated confidence intervals and effect sizes rather than the conventional p value threshold, finally, to confirm our findings we also implemented various statistical tools such as AIC, power analysis and bayes factor, statistical analysis was guided by^124–127^. For statistical analysis of data from the live imaging high content analysis experiments, data was normalized to logarithmic scale to achieve normally distributed features for statistical testing, sample size was computed to achieve 85% statistical power and the effect size (mean difference) 95% confidence intervals were computed for each feature from the selected features list using R. Following the analysis, Cohen’s effect size was computed for each feature using the statistical parameters obtained from the experiment itself in addition to the power of the experiment^128,129^. For Bayesians analysis used in live imaging high content analysis experiments the Bayes Factor was computed for each feature using the bayes package in R^130^. For high content IF imaging experiments, sample size was computed in order to achieve 85% statistical power using the pwr package in R^128,129^. The effect size (mean difference) confidence interval plots were computed using a bootstrap technique by using the dabstr package in R^131^. For the image-based acetylation experiment a One-Way ANOVA with Sidak’s correction for multiple comparisons was performed to compute p values. For the RNA-seq differential expression analysis, DESeq2^107^. Was used to compare GSD1a and HC groups. p values were corrected for multiple hypothesis testing using FDR correction. GSEA was performed on genes annotated to the binding sites using the R package, clusterProfiler and corrected for multiple hypothesis testing using FDR correction ^132,133^. For Metabolomics analysis to compare groups two-tailed t-tests were performed and raw p values were used to define significant metabolites, for MSEA raw p values were used as well. For the ATAC-seq affinity analysis of differential binding sites, DiffBind was used to compare GSD1a and HC groups. p values were corrected for multiple hypothesis testing using Benjamini Hochberg correction. GSEA was performed on genes annotated to the binding sites using the R package, clusterProfiler and corrected for multiple hypothesis testing using Benjamini Hochberg correction^132,133^. For the differential methylation of probes analysis performed on the EPIC methylation array data, the statistical model was built using limma^134^ and p values were corrected for multiple hypothesis testing using Benjamini Hochberg correction. For the differential methylation region analysis, a statistical model was built similarly and analyzed using DMRcate^135,136^. P values were corrected for multiple hypothesis testing using FDR correction. For the GSEA,methylGSA package was used^137^ based on genes annotated to the probes from the array, p values were corrected for multiple hypothesis testing using FDR correction. For the integration of ATAC-seq and EPIC data and RNA-seq permutations testing was done using R. Bootstrap correlation analysis was made using the boot package in R (1000 bootstraps per correlation analysis) and based on Pearson correlation and bootstrap mediated confidence interval around the correlation coefficients^138,139^. Enrichment analysis was performed using Enrichr^140–142^ and compared to the Jensen disease database. Multiple hypothesis correction was done using FDR.

#### Data analysis and integration analysis

Illustrations of experiments shown in the MS were created with <u>BioRender.com</u>. Generally, all data processing stages in all experiments were performed using Python 3 and R (vs 4.2.2), scripts are available in the link below. High quality image-based analysis is important and effective in biological data^143,144^. Specifically, in this work, prime Image analysis was customized for each experiment using the Harmony 4.8 image-analysis software. For live imaging high content analysis experiments analysis included quality control per plate, batch effect analysis following^145^ testing for unwanted sources of variation(i.e-gender, week of experiment) using variance calculation using the variance Partition package^146^ and as was shown by us^147^, well removal based on cell count, outlier identification^148^ to remove wells with extreme values and feature selection using backward feature elimination to remove correlated features and to select equal number of features per organelle to allow an unbiased approach for the multivariate analysis (See Figure S1a and Supplementary Table 1). To this end-features types (i.e-Texture, Area, Intensity) are not equal between different labeled organelles as a result of image analysis which generated feature types for each organelle based on segmentation preferences and biological interest. Finally, multivariate analysis highlighting ideal dimension reduction algorithms were chosen based on their overall fit to the data^149^. For RNA-seq data PCA was computed and plotted using the R package PCAtools^150^ Differential analysis visualization plots were executed using ggplot2^151^ and GSEA visualizations were done using clusterprofiler. For Metabolomics visualization of metabolites, s-PLS-DA analysis (based on 2 components) and MSEA analysis and visualization was performed and plotted using Metaboanalyst 5.0^109^. For ATAC-seq occupancy analysis was performed using DiffBind, PCA was computed and plotted using the R package PCAtools^150^. Affinity analysis visualization plots were executed using DiffBind and GSEA visualizations were done using clusterprofiler. Peaks were plotted using IGV and GVIZ^152^. For EPIC methylation analysis, K-means clustering analysis was performed using the R package^153^. PCA was computed using PCAtools^150^ and regression analysis was computed in R. For the integration of ATAC-seq, EPIC data and RNA-seq, data integration was guided and performed following^154,155^. s-PLS-DA was performed in R using mixOmics^156^ based on common genes shared across all datasets and normalized using z-scores, Heatmap for top genes was made using the R package ComplexHeatmap^157^, Venn diagrams of dataset contributions were made using ggplot2 amd ggvenndiagram^151,158^. Chromosome pie charts were made using ggplot2 and ggvenndiagram ^151,158^ based on significant genes from each data set (FDR<0.1). For the overlap analysis, significant genes were chosen based on significance in each dataset and represented by their most differential annotation region; genes from all datasets were then directed to the overlap analysis venn diagrams and their related plots were created using R, ggplot2 and ggVennDiagram packages^151,154,158^. For the bootstrap correlation analysis common genes shared across all datasets and normalized using z-scores were used, the chosen genes were then subjected to analysis in STRING^51^ K-means analysis using auto k clusters was performed to generate gene clusters tested for enrichment against all available databases in STRING. For the over representation enrichment analysis, genes were selected from the overlapping genes data set found in the overlapping analysis. Visualization was made using Plotly^159,160^. For biochemical and live imaging summary integration analysis, features were averaged and normalized to the same scale by z-scores to allow multi-level feature data to be integrated. Correlation analysis was performed in Python using pearson test^161^. Network analysis and K-means analysis^162^ were performed using factoextra on R. Visualization for several plots created throughout the manuscript were generated using Plotly^159,160^.

## Data and code Availability

Scripts for the main analysis sequences described could be found here - https://github.com/Urisprecher/GSD1a_MS_2024

Additional code and data are available upon request.

---

**Supplementary table 1-** Image-based HCA live imaging stats.

**Supplementary table 2-** Biochemical integration data.

**Supplementary table 3 -** RNA-seq Differential analysis list.

**Supplementary table 4 -** RNA-seq genes and metabolic genes from database.

**Supplementary table 5 -** Significant t-test metabolites from cell metabolomics analysis.

**Supplementary table 6 -** Significant t-test metabolites from media metabolomics analysis.

**Supplementary table 7 -** ATAC-seq Differential analysis list.

**Supplementary table 8 -** EPIC Differential probe analysis list.

**Supplementary table 9 -** EPIC Differential regions analysis list.

**Supplementary table 10 -** OMICS overlapping genes.

**Supplementary table 11 -** OMICS integration correlation genes.

**Supplementary table 12 -** EPIC bacr qc results.

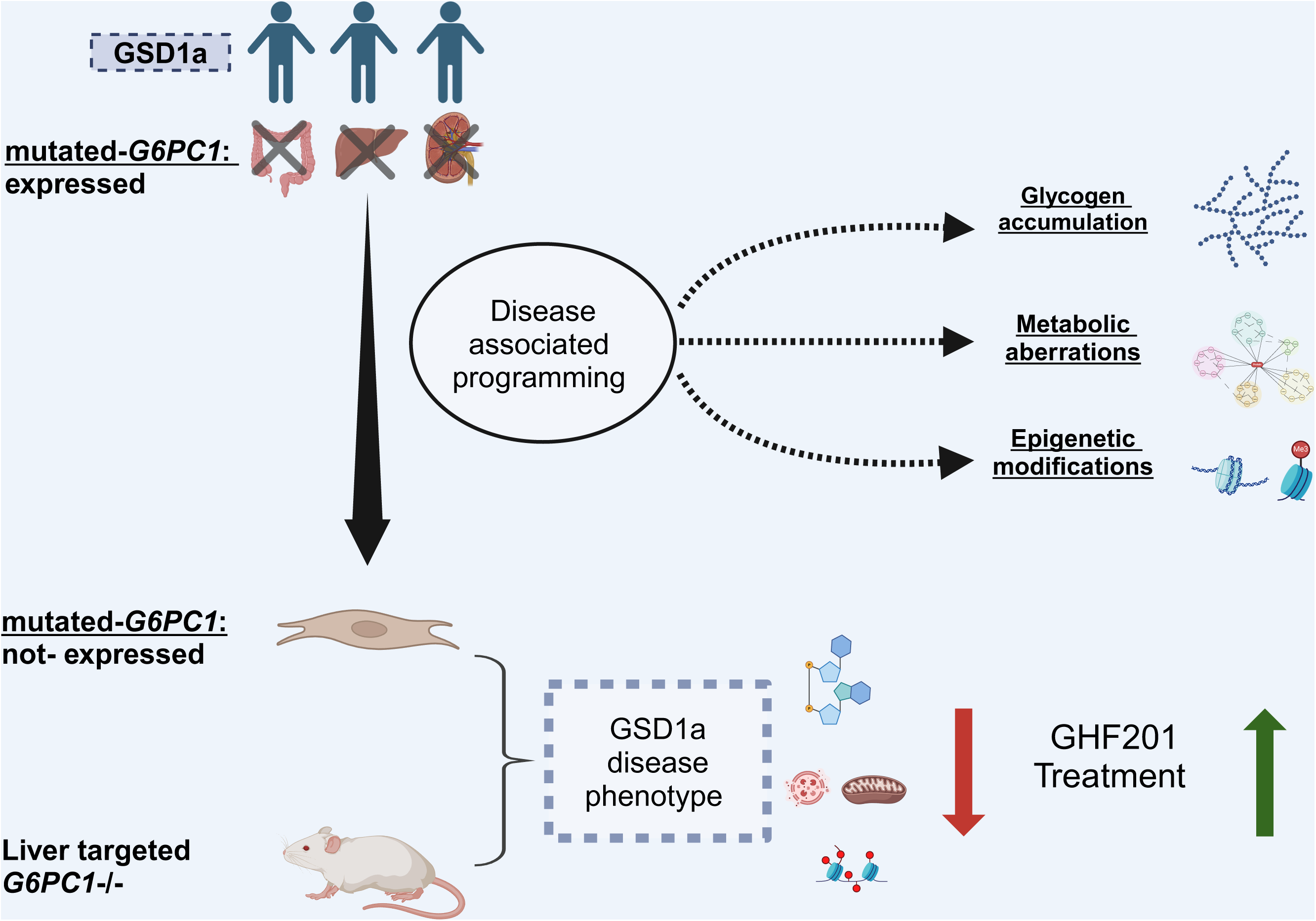

