## Supplementary Figures & Legends for "Disease-associated programming of cell memory in glycogen storage disorder type 1a"

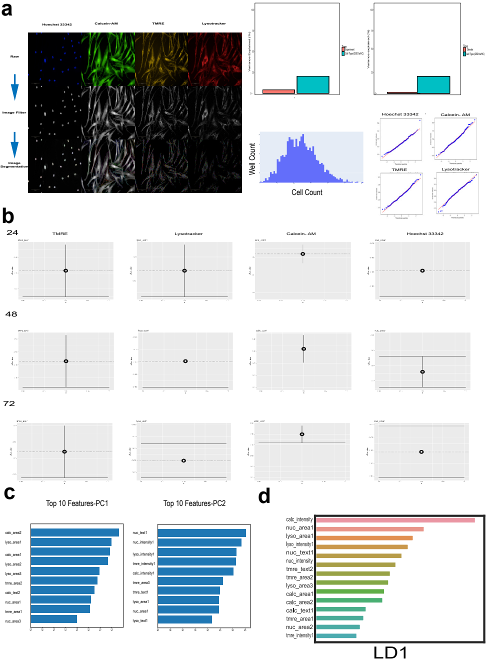

a
Hoechst 33342
Calcein-AM
TMRE
Lysotracker
Raw
Image Filter
Image Segmentation
Well Count
Cell Count
Hoechst 33342
Calcein- AM
TMRE
Lysotracker
b
24
TMRE
Lysotracker
Calcein- AM
Hoechst 33342
48
72
c
Top 10 Features-PC1
calc_area2
lyso_area1
calc_area1
lyso_area2
lyso_area3
tmre_area2
calc_text2
nuc_area1
tmre_area1
nuc_area3
Top 10 Features-PC2
nuc_text1
nuc_intensity1
lyso_intensity1
tmre_intensity1
calc_intensity1
tmre_area3
tmre_text1
lyso_area1
nuc_area1
lyso_text1
d
calc_intensity
nuc_area1
lyso_area1
lyso_intensity1
nuc_text1
nuc_intensity
tmre_text2
tmre_area2
lyso_area3
calc_area1
calc_area2
calc_text1
tmre_area1
nuc_area2
tmre_intensity1
LD1

**Supplementary Figure 1**

**a.** Left- representative raw, filtered and segmented images of GSD1a fibroblasts, showcasing the organelles labeled in the image-based HCA experiment using Hoechst 33342 for nucleus (blue), Calcein-AM for cytoplasm (green), TMRE for active mitochondria (yellow) and Lysotracker for lysosomes (red). Right, up- bar plots of explained variance in the image-based HCA experiment generated from the batch effect analysis for the following parameters respectively- experiment vs sample and gender vs sample. Bottom- histogram of cell count from wells analyzed in the image-based HCA experiment and representative QQ plots for one feature of each labeled organelle, indicating normality assessment.

**b.** Confidence Interval plots presenting representative features for each organelle in each time condition, from left to right - TMRE texture feature, Lysotracker texture feature, Calcein-AM texture feature and Hoechst 33342 area feature, rows show the condition from top to bottom, 24 h, 48 h, 72 h. N represents the number of wells analyzed for each group as shown in box plots in Figure 1. see Supplementary Table 1 for the description of each group.

**c.** The top ten contributing features to the variance explained by the first and second component of the PCA, including TMRE, Lysotracker, Calcein-AM and Hoechst 33342 features. (Features full names in Supplementary Table 1).

**d.** The top ten contributing features to the variance explained by the first component of the LDA, including TMRE, Lysotracker, Calcein-AM and Hoechst 33342 features. (Features full names in Supplementary table 1).

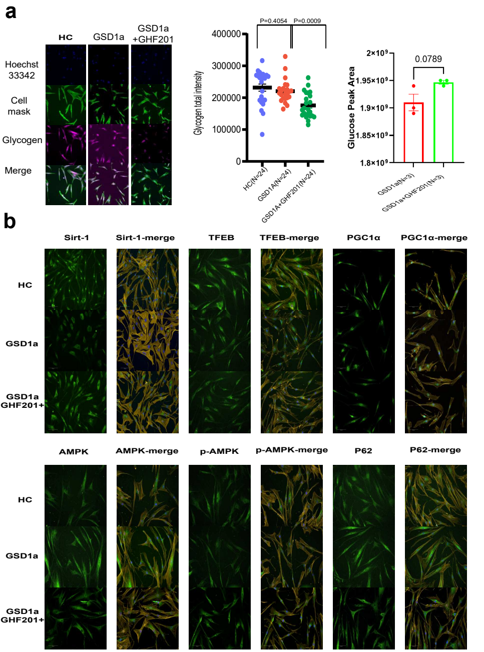
a
HC
GSD1a
GSD1a
+GHF201
Hoechst
33342
Cell
mask
Glycogen
Merge
P=0.4054
P=0.0009
Glycogen total intensity
400000
300000
200000
100000
0
HC(N=24)
GSD1A(N=24)
GSD1A+GHF201(N=24)
Glucose Peak Area
2×10⁹
1.95×10⁹
1.9×10⁹
1.85×10⁹
1.8×10⁹
0.0789
GSD1a(N=3)
GSD1a+GHF201(N=3)
b
Sirt-1
Sirt-1-merge
TFEB
TFEB-merge
PGC1α
PGC1α-merge
HC
GSD1a
GSD1a
GHF201+
AMPK
AMPK-merge
p-AMPK
p-AMPK-merge
P62
P62-merge
HC
GSD1a
GSD1a
GHF201+

**Supplementary Figure 2**

**a.** Left- representative images of a PAS assay experiment labeled organelles- Hoechst 33342 for nucleus (blue), Cell mask for cytoplasm (green) and Glycogen (magenta), comparing HC (blue) GSD1a(red) and GSD1a treated with 50 µM GHF201(green) fibroblasts. Middle- shown are scatter plots representing glycogen intensity, comparing intensity levels between HC (blue) GSD1a(red) and GSD1a with 50 µM GHF201 (green) fibroblasts (N represents the number of wells analyzed for each group, p values were imputed using two tailed t-tests). Right- Bar plots of glucose levels, determined by mass spectrometry, comparing GSD1a(red) and GSD1a treated with 50 µM GHF201(green) (N represents the number of samples analyzed for each group, p values were imputed using two tailed t-tests).

**b.** A panel of images obtained from the IF experiments performed throughout the manuscript, showing representative images for HC, GSD1a and GSD1a treated with 50 µM GHF201 fibroblasts from left to right. Markers are ordered from top to bottom: Sirt-1, TFEB, PGC1α, AMPK, p-AMPK. P62.

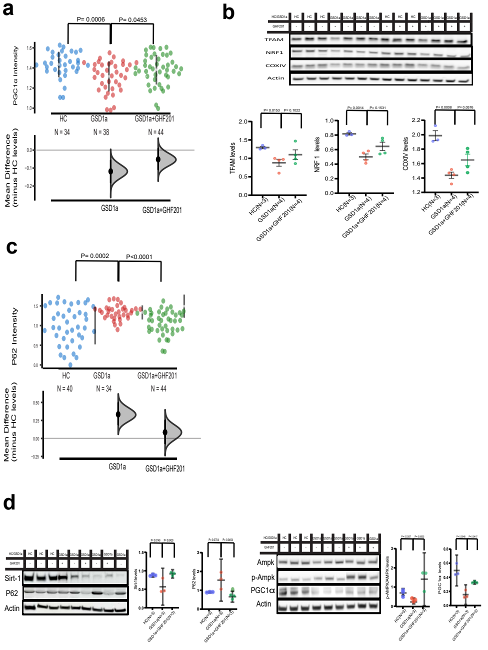
a
P= 0.0006
P= 0.0453
PGC1α Intensity
HC
GSD1a
GSD1a+GHF201
N = 34
N = 38
N = 44
Mean Difference (minus HC levels)
GSD1a
GSD1a+GHF201
b
TFAM
NRF1
COXIV
Actin
TFAM levels
NRF 1 levels
COXIV levels
HC(N=3)
GSD1a(N=4)
GSD1a+GHF201(N=4)
c
P= 0.0002
P<0.0001
P62 Intensity
HC
GSD1a
GSD1a+GHF201
N = 40
N = 34
N = 44
Mean Difference (minus HC levels)
GSD1a
GSD1a+GHF201
d
Sirt-1
P62
Actin
Ampk
p-Ampk
PGC1α
Actin

**Supplementary Figure 3**

**a.** Bootstrap CI Scatter plots representing PGC1α intensity normalized to HC levels. Comparative intensity levels between HC (blue), GSD1a (red) and GSD1a treated with 50 μM GHF201 (green) fibroblasts, on the right-hand side of each group a bar represents the mean intensity value with a 95% confidence interval. On the bottom the y-axis represents mean differences (effect size) between the tested groups to HC. A 95% confidence interval derived from a bootstrap resampling test is represented as a point estimate with a vertical bar and its respective distribution for each group effect size compared to HC (N indicates number of wells analyzed for each group, p values were computed using two-tailed t tests).

**b.** WB analysis of mitochondrial biogenesis and activity markers showing representative immunoblots (up) and densitometric quantifications (below) of comparative levels of TFAM, NRF1 and COXIV between HC (blue), GSD1a (red) and GSD1a treated with 50 μM GHF201(green) fibroblasts. Actin levels were used to normalize loading protein content of samples (N indicates samples analyzed for each group, p values were computed using two-tailed t tests).

**c.** Bootstrap CI Scatter plots representing P62 intensity normalized to HC levels. Comparative intensity levels between HC (blue), GSD1a (red) and GSD1a treated with 50 μM GHF201 (green) fibroblasts, on the right-hand side of each group a bar represents the mean intensity value with a 95% confidence interval. On the bottom the y-axis represents mean differences (effect size) between the tested groups to HC. A 95% confidence interval derived from a bootstrap resampling test is represented as a point estimate with a vertical bar and its respective distribution for each group effect size compared to HC (N indicates the number of wells analyzed for each group, p values were computed using two-tailed t tests).

**d.** WB analysis of Sirt-1, P62, P-AMPK, AMPK and PGC1α (from left to right) showing representative immunoblots (up) and densitometric quantifications (to the right) of the comparative levels of each protein between HC (blue), GSD1a (red) and GSD1a treated with 50 μM GHF201(green) fibroblasts. Actin levels were used to normalize loading protein content of samples (N indicates samples analyzed for each group, p values were computed using two-tailed t tests).

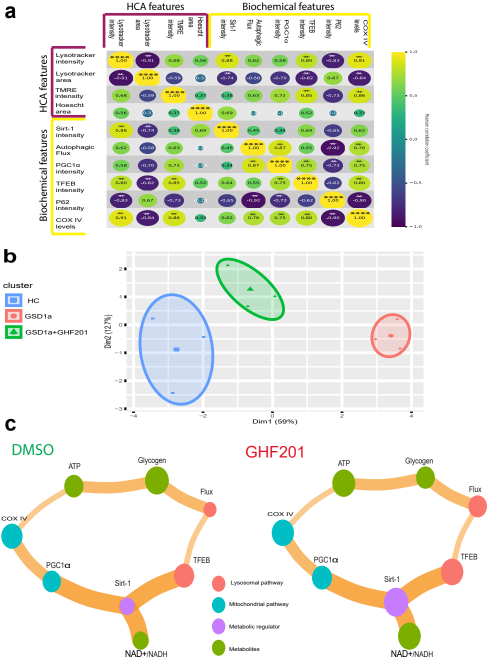
a
HCA features
Biochemical features
Lysotracker intensity
Lysotracker area
TMRE intensity
Hoescht area
Sirt-1 intensity
Autophagic Flux
PGC1α intensity
TFEB intensity
P62 intensity
COX IV levels
Pearson correlation coefficient
1.0
0.5
0.0
−0.5
−1.0
b
cluster
HC
GSD1a
GSD1a+GHF201
Dim2 (12.7%)
Dim1 (59%)
c
DMSO
GHF201
ATP
Glycogen
Flux
COX IV
PGC1α
Sirt-1
TFEB
NAD+/NADH
Lysosomal pathway
Mitochondrial pathway
Metabolic regulator
Metabolites

**Supplementary Figure 4**

**a.** Correlation bubble plot representing a Pearson correlation analysis performed on GSD1a and HC samples, measuring the interaction between selected features from the image-based HCA experiment and several biochemical derived markers assessed in Figures 1-5. Features are categorized as HCA or biochemical-analysis based. The Radius and color of the circles reflects the correlation coefficient value; significant correlations are noted by asterisks (N=3 per group, significant p values <0.1).

**b.** K-means clustering plot calculated by K-means algorithm (k=3) using data from HC (blue), GSD1a (red) and GSD1a treated with 50 µM GHF201 fibroblasts. Integrating features obtained from several biochemical derived markers assessed in Figures 1-5, (N=3 per group) The circle of each cluster reflects a 95% confidence interval.

**c.** Network analysis visualization presents the interaction between key markers in GSD1a samples (left) and GSD1a treated with 50 µM GHF201 fibroblasts(right)( N=3 per group). The node color indicates the marker pathway, node size represents the measured value of the marker. Edge weight indicates the strength of the marker’s interaction based on the literature.

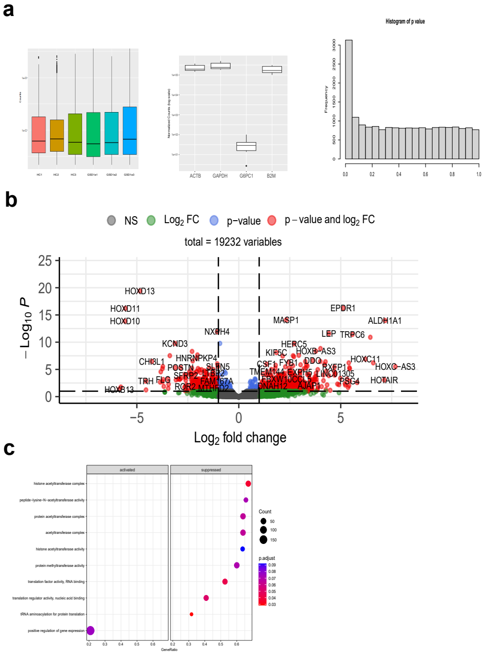
a
HC1
HC2
HC3
GSD1a1
GSD1a2
GSD1a3
ACTB
GAPDH
G6PC1
B2M
Histogram of p value
Frequency
b
NS
Log2 FC
p−value
p − value and log2 FC
total = 19232 variables
− Log10 P
Log2 fold change
HOXD13
HOXD11
HOXD10
EPDR1
MASP1
ALDH1A1
NXPH4
LEP
TRPC6
KCND3
HERC5
HOXB−AS3
CHI3L1
HNRNPKP4
KIF5C
HOXC11
POSTN
SLFN5
CSF1
FYB1
DDO
RXFP1
HOXC−AS3
SFRP2
LTBR2
TMEM144
EXPH5
LINC01305
TRH
FLG
FAM167A
FBXW10
CCL5
PSG4
HOTAIR
HOXB13
ROR2
MTHFD2
DNAH12
AJAP1
c
activated
suppressed
histone acetyltransferase complex
peptide−lysine−N−acetyltransferase activity
protein acetyltransferase complex
acetyltransferase complex
histone acetyltransferase activity
protein methyltransferase activity
translation factor activity, RNA binding
translation regulator activity, nucleic acid binding
tRNA aminoacylation for protein translation
positive regulation of gene expression
Count
p.adjust
GeneRatio

**Supplementary Figure 5**

**a.** Left- box plots of gene counts(y axis) from 3 HC and 3 GSD1a fibroblasts (x axis) included in the RNA-seq experiment. Middle- box plots of normalized gene counts of housekeeping genes- ACTB, GAPDH and B2M compared to G6PC1 across all samples. Right- p value histogram generated from DESEQ2 differential analysis on RNA-seq data comparing HC and GSD1a samples.

**b.** Volcano plot generated from DESEQ2 differential analysis and comparing HC and GSD1a samples on ~19000 genes that passed RNA-seq QC steps( FDR cutoff =0.1, log 2 fold change cutoff =1, N =3 and represents samples analyzed).

**c.** Dot plot of GSEA GO pathways (activated or suppressed) based on genes from the RNA-seq experiment and comparing HC to GSD1a samples. Adjusted p values were generated using FDR and are represented by the color gradient shown next to the plot. Size of the dots reflect gene counts per pathway.

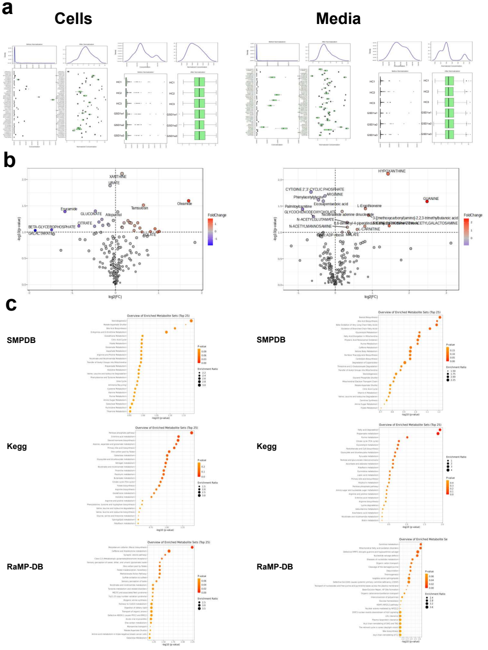
a
Cells
Media
b
XANTHINE
URATE
Erucamide
GLUCONATE
Allopurinol
Tamsulosin
Oleamide
BETA-GLYCEROPHOSPHATE
GALACTARATE
CITRATE
FoldChange
log2(FC)
-log10(p-value)
HYPOXANTHINE
CYTIDINE 2',3'-CYCLIC PHOSPHATE
ARGININE
Phenylacetylglycine
GUANINE
Eicosapentaenoic acid
L-Ergothioneine
Palmitoylcarnitine
GLYCOCHENODEOXYCHOLATE
N-ACETYLGLUTAMATE
N-ACETYLMANNOSAMINE
L-CARNITINE
MALATE
c
SMPDB
Kegg
RaMP-DB
Overview of Enriched Metabolite Sets (Top 25)
P-value
Enrichment Ratio
-log10 (p-value)

**Supplementary Figure 6**

**a.** Left- histograms and box plots of pre-normalzation and post-normalization of features (metabolites) and samples ( 3 HC and 3 GSD1a fibroblasts) from the cell metabolomics experiment. Right- histograms and box plots of pre-normalzation and post-normalization of features (metabolites) and samples( 3 HC and 3 GSD1a fibroblasts) from the media metabolomics experiment.

**b.** Left- Volcano plots from the cell metabolomics experiment. Right- Volcano plots from the media metabolomics experiment. Fold change levels are shown as a color gradient, significant metabolites (p value<0.1) are noted.

**c.** Left- Dot plots based on MSEA against SMPDB, KEGG and RaMP-DB ( top to bottom) pathways performed on cell metabolomics and comparing HC to GSD1a samples, raw p values are represented by the color gradient shown next to the plot. Size of the dots reflect enrichment ratio per pathway. Right- Dot plots based on MSEA against SMPDB, KEGG and RaMP-DB ( top to bottom) pathways performed on media metabolomics and comparing HC to GSD1a samples, raw p values are represented by the color gradient shown next to the plot. Size of the dots reflect enrichment ratio per pathway.

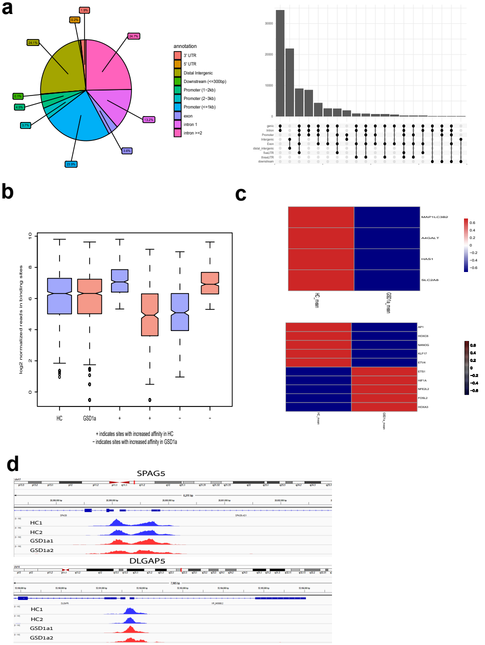
a
1.9%
0.2%
24.7%
24.1%
0.1%
4.5%
3.7%
13.2%
3.6%
23.9%
annotation
3' UTR
5' UTR
Distal Intergenic
Downstream (<=300bp)
Promoter (1−2kb)
Promoter (2−3kb)
Promoter (<=1kb)
exon
intron 1
intron >=2
30000
20000
10000
0
genic
intron
Promoter
Intergenic
Exon
distal_intergenic
fiveUTR
threeUTR
downstream
b
log2 normalized reads in binding sites
HC
GSD1a
+
+
−
−
+ indicates sites with increased affinity in HC
− indicates sites with increased affinity in GSD1a
c
MAP1LC3B2
A4GALT
HAS1
SLC2A8
HC_mean
GSD1a_mean
AP1
HOXC6
NANOG
KLF17
ETV4
ETS1
HIF1A
NFE2L2
FOSL2
HOXA3
d
SPAG5
HC1
HC2
GSD1a1
GSD1a2
DLGAP5
HC1
HC2
GSD1a1
GSD1a2

**Supplementary Figure 7**

**a.** Left- A pie chart of the distribution of annotated regions in the analyzed peaks generated from the ATAC-seq experiment. Right- an upset plot presenting the annotated regions in the analyzed peaks generated from the ATAC-seq experiment

**b.** Left- Box plots of normalized log2fc of reads of binding sites for the groups shown as increased (+) or decreased (-) affinity binding sites, HC box plots are shown in blue and GSD1a boxplot are shown in red.

**c.** Heatmap of expression levels between HC and GSD1a genes shown in the motif analysis.

**d.** IGV plots from the ATAC-seq experiment of two control genes which are highly expressed in fibroblasts (SPAG5 and DLGAP5) showing the peaks of these genes are high in all samples analyzed for HC ( blue) and GSD1a(red) fibroblasts.

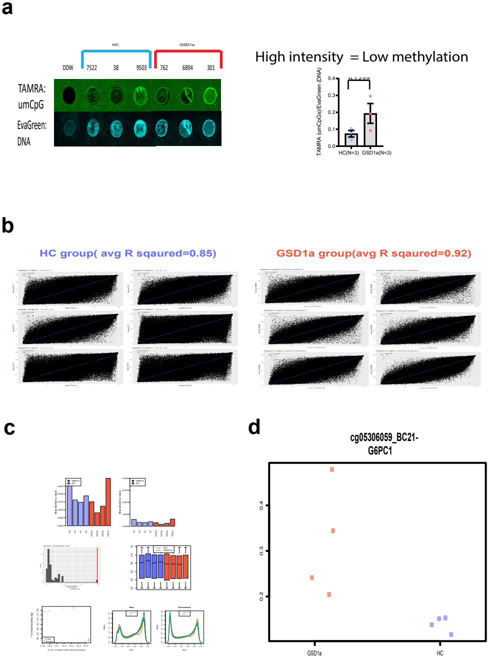
a
HC
GSD1a
DDW
7522
38
9503
762
6894
301
TAMRA: umCpG
EvaGreen: DNA
High intensity = Low methylation
TAMRA (umCpGs)/EvaGreen (DNA)
0.1232
HC(N=3)
GSD1a(N=3)
b
HC group( avg R sqaured=0.85)
GSD1a group(avg R sqaured=0.92)
c
d
cg05306059_BC21-
G6PC1
GSD1a
HC

**Supplementary Figure 8**

**a.** Left- Microarray slide of DNA from HC and GSD1a fibroblasts, selectively marked by EvaGreen (DNA dye- down) and TAMRA (unmethylated CpG regions- up). Right- quantification of the above-mentioned analysis comparing unmethylated CpG regions levels normalized to total DNA between HC (blue) and GSD1a (red) fibroblasts (N represents samples analyzed for each, p values were computed using two tailed t-tests).

**b.** Scatter plots representing all probes analyzed for each sample in the EPIC experiment. Each scatter plot presents two HC or two GSD1a samples against each other and the R coefficient of a linear model. Left- scatter plots of the HC samples, Right- scatter plots of the GSD1a samples, On the top -average R coefficients for each group.

**c.** Processing and quality control plots generated from the EPIC array analysis pipeline. Up- p values bar plots showcasing all samples validity to be analyzed. Middle- Quantro analysis plots. Bottom left - sex identification plots. Bottom right- raw and normalized M values generated for the analysis.

**d.** A scatter plot showing the different methylation levels measured between HC(blue) and GSD1a (red) fibroblasts on the G6PC1 gene CpG probe (N represents samples analyzed for each group=4).

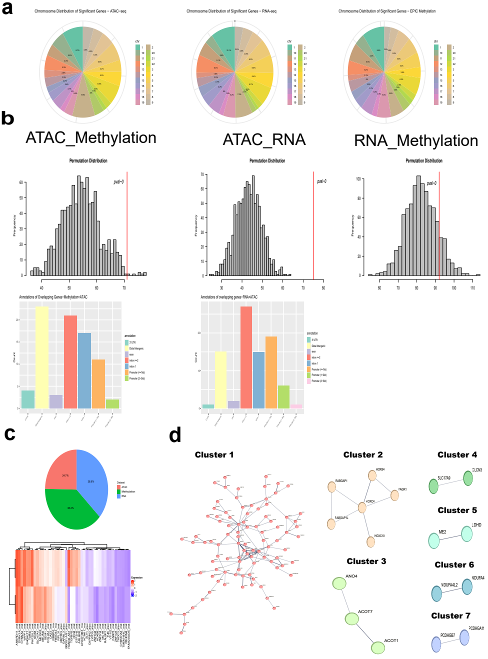
a
Chromosome Distribution of Significant Genes ~ ATAC-seq
Chromosome Distribution of Significant Genes ~ RNA-seq
Chromosome Distribution of Significant Genes ~ EPIC Methylation
b
ATAC_Methylation
ATAC_RNA
RNA_Methylation
Permutation Distribution
pval~0
Frequency
Annotations of Overlapping Genes~Methylation+ATAC
Annotations of overlapping genes~RNA+ATAC
c
d
Cluster 1
Cluster 2
Cluster 3
Cluster 4
Cluster 5
Cluster 6
Cluster 7
RABGAP1
HOXB4
PAGR1
HOXC4
RABGAP1L
HOXC10
ANO4
ACOT7
ACOT1
SLC17A9
CLCN3
ME2
LDHD
NDUFA4L2
NDUFA4
PCDHGB7
PCDHGA11

**Supplementary Figure 9**

**a.** Pie charts of the distribution of chromosomes of significant genes coming from ATAC-seq ( left), RNA-seq(middle) and EPIC methylation (right) datasets.

**b.** Histograms of permutation of overlapping significant genes generated from the overlap analysis of ATAC-seq and EPIC methylation (left), ATAC-seq and RNA-seq(middle) and RNA-seq and EPIC methylation(right), p value of permutation is indicated in the plot. Underneath the histogram of ATAC-seq and EPIC methylation and ATAC-seq and RNA-seq a bar plot with the annotations of the overlapping genes between the indicated datasets.

**c.** Up- pie chart of the distribution EPIC methylation and ATAC-seq and RNA-seq gene contribution to sparse-pls-DA plot in Figure 8d. Down- Heatmap of the top contributing genes for sparse-pls-DA analysis coming from different datasets, Fold change is indicated as a scale bar.

**d.** STRING k means analysis clusters of the bootstrap correlation analysis.

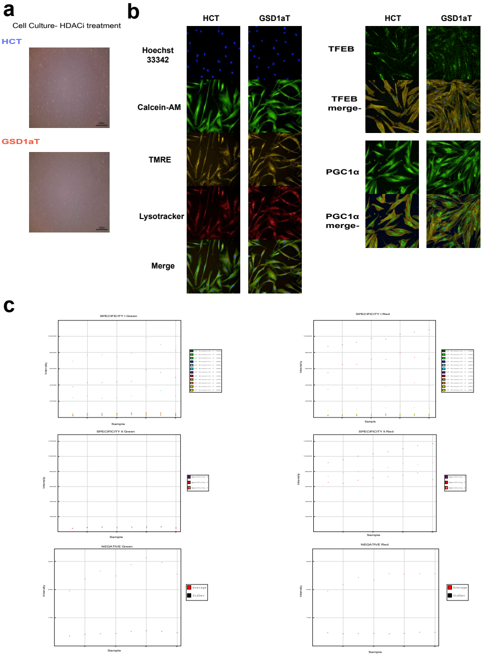
a
Cell Culture- HDACi treatment
HCT
GSD1aT
b
HCT
GSD1aT
Hoechst 33342
Calcein-AM
TMRE
Lysotracker
Merge
TFEB
TFEB merge-
PGC1α
PGC1α merge-
c
SPECIFICITY I Green
SPECIFICITY I Red
SPECIFICITY II Green
SPECIFICITY II Red
NEGATIVE Green
NEGATIVE Red
Intensity
Sample
Average
StdDev

**Supplementary Figure 10**

**a.** Cell Culture images taken using an Evos microscope indicating the quality of cells following HDACi treatment.

**b.** Representative images obtained from the IF and HCA experiments performed on HC and GSD1a cells treated with HDACi. For the HCA panel, from top to bottom - Hoechst 33342 for nucleus (Blue), Calcein-AM for cytoplasm (Green), TMRE for active mitochondria (Yellow) and Lysotracker for lysosomes (Red). For IF, from top to bottom- TFEB and PGC1a.

**c.** Representative plots indicating the quality control metrics of the illumina EPIC methylation array.
